# Non-invasive imaging of gene expression and protein secretion dynamics in living mice: identification of ectopic prothrombin expression as driver of thrombosis in cancer

**DOI:** 10.1101/2021.07.08.451623

**Authors:** Jamie Nourse, Sergey Tokalov, Shazad Khokhar, Essak Khan, Lina K. Schott, Lisa Hinz, Ludwig Eder, Danielle Arnold-Schild, Johanna Satrapa, Karl J. Lackner, Hugo Ten Cate, Hans Christian Probst, Sven Danckwardt

**Affiliations:** Posttranscriptional Gene Regulation, University Medical Centre Mainz, Germany; Institute for Clinical Chemistry and Laboratory Medicine, University Medical Centre Mainz, Germany; Centre for Thrombosis and Hemostasis (CTH), University Medical Centre Mainz, Germany; Research Center, Immunology, University Medical Centre Mainz, Germany; Thrombosis Expertise Center, Department of Internal Medicine, Maastricht University Medical Center and Cardiovascular Research Institute Maastricht (CARIM), Maastricht, the Netherlands; German Centre for Cardiovascular Research (DZHK), Berlin, Germany; Centre for Healthy Aging (CHA), Mainz, Germany

**Keywords:** Prothrombin, liver, blood coagulation, secretory protein, cancer, thromboembolism, in vivo imaging, luminescence

## Abstract

The topology of gene expression and protein localization is a crucial characteristic of life, where the spatiotemporal dynamic of secretory proteins instruct higher order organization, including the orchestration of developmental and adaptive programs. However tools to non-invasively interrogate the fate of secretory proteins in vivo are scarce. Here we introduce a genetic tagging strategy for in vivo imaging of the secretion and expression dynamics of secretory proteins in living animals. Applying this to a prototypical liver-derived secretory protein, we demonstrate that this approach, combined with optical in-vivo imaging, uncovers extrahepatic prothrombin expression in multiple novel anatomical sites (including testes, placenta, brain, kidney, heart and lymphatic system) and in emerging tumors, resulting in significant amounts of tumor-derived prothrombin in the blood with procoagulant properties. Syngeneic cell lines from this mouse model enable unravelling regulatory mechanisms in high resolution, and in a scalable format ex vivo. Beyond discovering new functions of proteins in a targeted manner, this model allows identifying rheostats in the cross-talk between gene expression and availability of a secretory protein. It is also a valuable resource for uncovering novel (tissue-specific) therapeutic vulnerabilities.

## Main

Cells synthesize thousands of proteins with diverse functions, which need to be directed to specific locations in accurate amounts at precise time^1^. Nearly one-third of the human proteome is targeted to secretory environments consisting of the endoplasmic reticulum, lysosome, plasma membrane or the extracellular space^2^ by a well-operating cargo system^3,4^. Almost all cytokines, hormones, receptors, peptidases, channels, extracellular matrix components, transport proteins and coagulation factors are clients of this machinery^5^. Secretory proteins also possess a central role in anatomical and functional compartmentalization, for example by controlling the function, volume and integrity of the vascular compartment^6-8^. Dysfunction of the production and proper delivery of secretory proteins is the cause of a variety of diseases, including neurodegenerative, developmental and cardiovascular disorders, and perturbations of the hemostatic system^9-14^.

In most cells, secreted proteins account for 10-20% of the transcriptome^2^. In contrast, about 40% of all transcripts in the liver encode secreted proteins, reflecting the particular functional demands of this metabolic organ. The liver synthesizes and releases into the bloodstream numerous secretory proteins, including albumin, the most abundant serum protein regulating the colloid osmotic pressure and serving as carrier protein, and most coagulation factors^15^. Accordingly, the impairment of hepatic protein production and secretion results in detrimental consequences^16^. This includes bleeding disorders and venous thromboembolism, a global cause of mortality^17^, exemplifying the importance of a well-balanced production of liver-derived secretory proteins^14^. Regulatory mechanisms have evolved to ensure the required functional levels of secretory proteins^18^ and to adapt production to conditions of increased turnover and demand^19^. This likely also involves hitherto largely undefined (auto) regulatory mechanisms reminiscent of hormone feedback circuits. Depending on spatial and temporal availability, secretory proteins can also display entirely distinct functional characteristics^20-22^. Illuminating these processes at the systems level paves the way to unravel underlying regulatory principles and disease-eliciting mechanisms. It can also help identify novel therapeutic opportunities^23^.

Molecular imaging is a rapidly emerging field, providing quantitative, non-invasive visual representations of fundamental biological processes in intact living subjects^24^. In addition to innovative new probes and imaging agents^25^, current applications of non-invasive imaging involve reporter genes to monitor processes including signal transduction, cell tracking, or transgene expression, with exciting advances employing synthetic biology coupled to optoacoustic imaging^26^, and innovative applications in humans^27^. Traditional non-invasive imaging of endogenous gene expression can be complex^28,29^. While indirect imaging relies on reporter gene expression, for example serving as a proxy for the abundance of transcription factors, direct measurements require the manipulation of the endogenous gene without impairing the epigenetic, transcriptional and posttranscriptional regulation^30-32^. Despite the critical medical dimension, tools to non-invasively visualize expression and protein secretion dynamics, and spatiotemporal distribution of secretory proteins at a systems level are surprisingly scarce.

Here we report on a novel multimodal reporter animal designed for conditional non-invasive optical imaging of a prototypical liver-derived secretory protein^33^. This is based on multicolor imaging to dissect the expression and secretion dynamics of prothrombin in its physiological context in a living animal. In a proof-of-concept, including the validation with an inducible hypomorphic animal, we confirm that the reporter mouse model faithfully recapitulates known modifiers of prothrombin expression. We also discover extrahepatic prothrombin expression at various anatomical sites, and in emerging malignant mesenchymal tumors, resulting in significant quantities of functional tumor-derived prothrombin in the plasma, which entertains a procoagulant function. It thereby documents the autonomous expression of a secretory protein that becomes harmful if aberrantly expressed. We further demonstrate that complementary syngeneic primary cell culture lines obtained from this reporter animal enable the deconvolution of regulatory mechanisms in high resolution, and in a scalable high throughput format ex vivo. It thereby represents a valuable resource for uncovering disease-eliciting cues and novel therapeutic vulnerabilities.

## Results

Thrombin is the key serine protease involved in blood coagulation and hemostasis. It is synthesized in the liver as an inactive 72-kDa precursor protein (prothrombin) and secreted into the blood circulation. Upon activation, thrombin converts fibrinogen into fibrin, the main component of blood clots, and activates platelets. Besides its role in hemostasis^34^, thrombin is involved in numerous other processes, including embryonic development, angiogenesis, wound healing, inflammation, atherosclerosis and tumor biology^35,36^. This reflects the broad action of thrombin on membrane-bound G-protein coupled protease activated receptors (PARs)^37^. However, many of these actions naturally affect cells residing in the extravascular compartment^34^. This questions the prevailing observation of prothrombin synthesis being confined to the liver^2^.

Here we set out to generate a multimodal imaging reporter animal to characterize the spatiotemporal dynamics of this pleiotropic liver-derived protein in vivo. Specifically, our strategy aimed at creating a reporter for (1) identifying and quantifying (non)canonical origins of prothrombin expression, and (2) disentangling gene expression and protein secretion dynamics under control of endogenous regulatory mechanisms. Further it ought to feature a (3) differential labeling of the secretory protein (depending on the origin where it is expressed and secreted), provide a (4) component to explore the downstream function(s) and serve as (5) resource for isogenic primary reporter cell lines for further in-depth studies, including regulatory pathways by high content analyses in a scalable format ex vivo.

To generate this versatile tool, we employed a multicistronic reporter strategy targeting the endogenous prothrombin (F2) gene locus in B6 mice (Fig. 1a). We generated a conditional knock-in allele for in-frame insertion of two luminescence reporters, Rluc8.6-535^38^ and Fluc2CP (with the latter harboring two protein destabilization domains, hCL1 and hPEST, for improved reporter dynamics), a bright and stable near-infrared fluorescent protein (iRFP) for in vivo imaging^39^ and a human Diphtheria-toxin receptor (DTR) directly upstream of the translation termination codon and the 3’ untranslated region. By using highly efficient viral P2A co-translational ‘self-cleaving’ peptides^40^, the endogenous prothrombin and the inserted reporters are co-expressed as individual proteins in a strict 1:1:1 stoichiometry^41^. While prothrombin is tagged by in-frame fusion to Rluc and secreted into the circulation, the Flag-iRFP, the DTR and the Fluc are cell-resident (and thereby ‘label’ cells that express prothrombin). For the downstream functional exploration of prothrombin and/or the cell(s) expressing this secretory protein, a BRET-compatible Rluc label was chosen (to allow for ligand-receptor interaction^42^ and compartment colocalization studies^43^) in addition to the DTR for targeted cell depletion with Diphtheria toxin (DTX)^44^. For a Cre-mediated excision^45^, two loxP-sites were inserted upstream of the Rluc and downstream of the DTR. This enables differential labeling of the canonical (liver-derived) and non-canonical (extrahepatic) prothrombin and a conditional DTX-mediated depletion of prothrombin-expressing cells.

**Figure 1.**
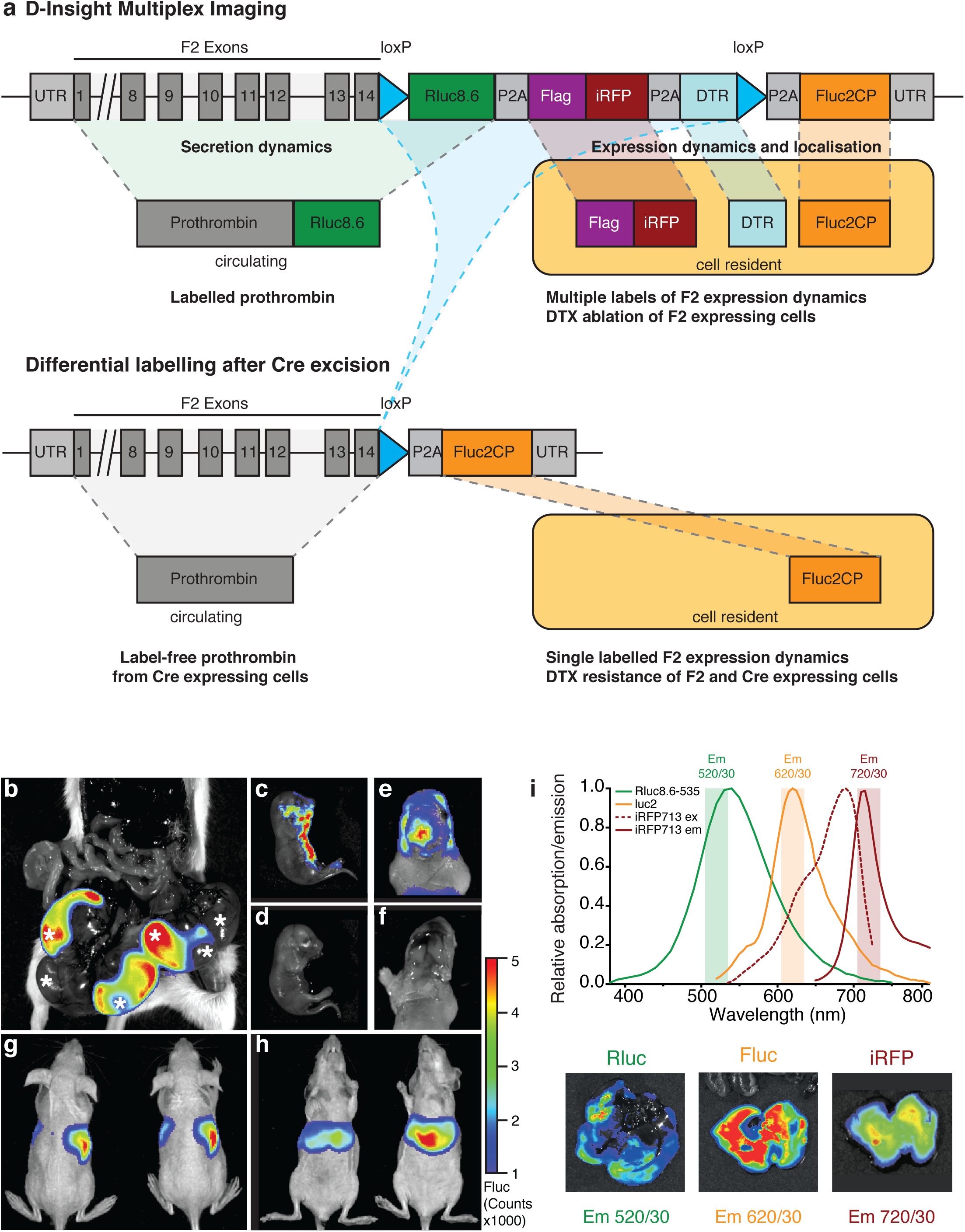
D-Insight mouse model for conditional real time *in vivo* imaging of the spatiotemporal dynamics of prothrombin (F2) gene expression and secretion. **a**, Schematic representation of the F2 D-Insight model design, carrying various labels to visualize F2 gene expression and secretion dynamics by non-invasive real-time in vivo (3D) imaging (Rluc8.6 = red shifted Renilla luciferase Rluc8.6-535; iRFP = near infrared fluorescent protein; DTR = Diphtheria Toxin Receptor; Fluc2CP = Firefly-Luciferase, P2A = co-translational ‘self-cleaving’ peptide, loxP = sequence for Cre-mediated recombination). **b-i**, Multiplex F2 expression labelling in vivo. **b**, Postmortem bioluminescence-imaging (Fluc 10 min after luciferin substrate i.p. injection) of a pregnant B6 animal (after pairing with heterozygous male D-Insight animal) showing 3 D-Insight positive, out of 6 embryos (each embryo is marked with an asterisk). **c**, D-Insight positive embryo (3 weeks old) showing F2 gene expression along the ventral side of the cranio-caudal axis and **e**, in the brain (same animal after craniotomy) compared to B6 litter control animals **d**,**f. g**-**h**, Hepatic expression in adult animals imaged dorsally and ventrally. **i**, spectra of multiplex labeling and imaging of liver with respective emission filters.

Despite the extensive genetic modifications, we obtained viable reporter animals in expected Mendelian ratios (Fig. 1b). The obtained Fluc signals are specific for the transgene, and in the developing embryo project to the ventral part of the cranio-caudal axis (Fig. 1c,d) and the brain (Fig. 1e,f). This corresponds to earlier reports supposing prothrombin expression at various locations in the developing embryo^46^ and a function in the nervous system^47^. In contrast, adult animals show strong signals (Fluc, Rluc and iRFP) that co-localize with the liver (Fig. 1g-i), corresponding to the hepatic expression of prothrombin in adults (www.proteinatlas.org).

To validate the functionality of the DTR introduced with our multicistronic transgene, we next administered DTX to the reporter and B6 controls animals. Expectedly, we observe that 24h after DTX injection the Fluc signal declines in the liver (Extended Data Fig. 1a,b), reflecting a DTX-mediated disintegration of this organ. This is mirrored by a significant induction of the transaminases (GOT, GPT), reflecting liver tissue damage (Extended Data Fig. 1c). In contrast, B6 animals that do not harbor the human DTR do not show an induction.

For most applications, the DTX-mediated ablation of prothrombin expressing cells (primarily targeting the liver) is incompatible with a functional assessment of minor sites of prothrombin expression. Additionally, we aimed to exploit a differential labeling strategy that allows us to distinguish prothrombin produced in the liver from extrahepatic sites (Fig. 1a). To validate the Cre-mediated conditional labeling, we crossed D-Insight animals with a liver-specific Cre-expressing line (Albumin-Cre) and analyzed the bioluminescence of the offspring. Expectedly, we obtained animals in which the Rluc fused in-frame to the prothrombin can no longer be detected, while Fluc expression remained intact (Extended Data Fig. 1d-g). This manipulation also results in the excision of the DTR (Fig. 1a), which enables a selective DTX-mediated ablation of prothrombin-expressing extrahepatic cells. The elimination of Rluc and DTR thereby introduces a functional component to further explore downstream features of extrahepatic prothrombin and/or the function of prothrombin-expressing cells in vivo.

Our reporter is designed to discriminate between prothrombin expression and secretion (Fig. 1a). Accordingly, while Rluc and Fluc can be detected in liver lysates, only the Rluc signal can be found in the plasma (Fig. 2a). Comparing plasma thrombin activities of B6 and D-Insight reporter animals revealed almost identical levels (Extended Data Fig. 2a), demonstrating that the Rluc tag per se does not impede protein transport nor thrombin function.

**Figure 2.**
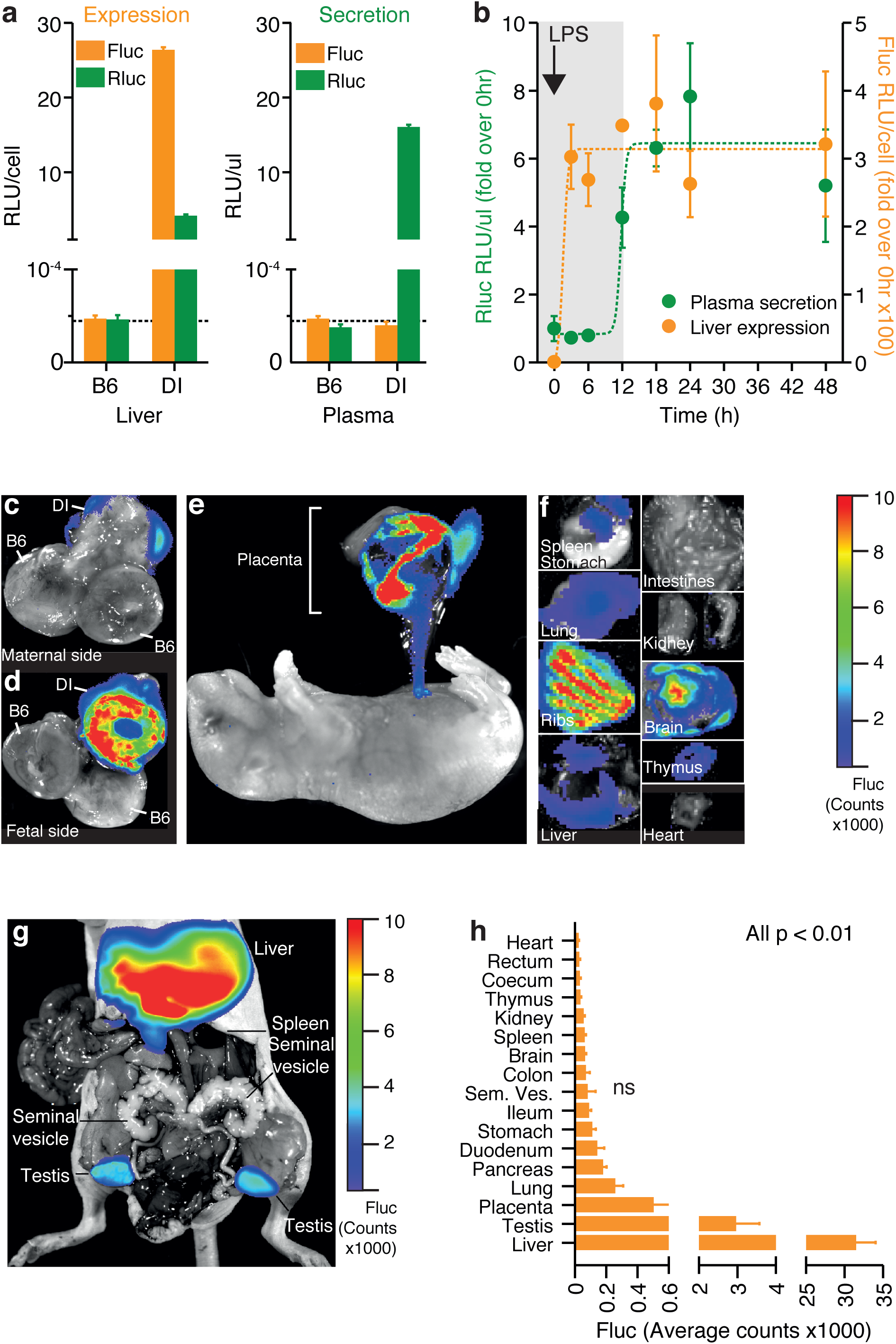
Differential labeling distinguishes expression and secretion dynamics and identifies numerous extrahepatic sources of prothrombin in embryos and adult animals. **a**, Labelled prothrombin is secreted into the circulation. While Rluc and Fluc can be detected in liver lysates, only Rluc (fused to prothrombin, Fig. 1a) is secreted and can thus be detected in the plasma. **b**, Expression and secretion dynamics in septicemic D-Insight animals (after LPS injection) showing early induction of F2 gene expression (Fluc in liver) with retarded accumulation of Rluc-tagged prothrombin in the blood circulation (Rluc in plasma). Non-linear regressions (Sigmoidal dose-response) are indicated by dashed lines. Established and novel extrahepatic expression of prothrombin (Fluc) in the embryo **c-f** (**c** and **d**, maternal and fetal part of placenta of a pregnant B6 animal after pairing with a heterozygous male D-Insight animal, in **e** the pseudocolor overlay shows reporter expression only for the placenta and umbilical cord) and in adults **g-h** (for simplicity, the baseline Fluc bioluminescence of reporter negative B6 controls animals, averaging to 1-2 Fluc counts, is not depicted in **h**; ns = not significant).

Next, we aimed to validate the reporter with established modifiers of prothrombin expression. Prothrombin is induced during septicaemia by a regulatory mechanism controlling RNA processing^19^. We injected lipopolysaccharides (LPS) into reporter animals and observed a significant upregulation of prothrombin expression (Fluc intensity) in the liver 3 hours after LPS-injection (Fig. 2b) - reminiscent of earlier reported RNA induction. While this induction is rapid, the abundance of prothrombin in plasma (Rluc) increases with retarded kinetics 12 hours after LPS-injection (Fig. 2b). This demonstrates that the reporter mouse model recapitulates established prothrombin expression dynamics^19,48^. By differential labeling and quantification of the expression and secretion dynamics, it allows disentangling these processes in vivo.

We next tested the accuracy and sensitivity of the reporter system, and whether new anatomical origins of this prototypic liver-derived protein can be detected. The knockout of prothrombin is embryonically lethal with phenotypes associated with impaired vascular integrity during development ^49,50^. Based on this, we expected to retrieve prothrombin expression characteristics that can be associated with these findings. We investigated prothrombin expression in developing embryos, and observed that the fetal part of the placenta (and the umbilical cord) expresses significant amounts (Fig. 2c-e). We also identified expression in a variety of other organs in the developing embryo (Fig. 2f). We further extended these studies to adult animals. We identified significant amounts of prothrombin expression in the testis (Fig. 2g) and at numerous previously unreported extrahepatic locations (Fig. 2h), including the lung, pancreas, lymphatic system (spleen), kidney and heart.

To further confirm the accuracy of the reporter system, we compared Fluc signals of various tissues from D-Insight animals and compared this to prothrombin RNA abundance (Extended Data Fig. 2b). This revealed that the reporter activity closely mirrors prothrombin gene expression over more than 2 orders of magnitude, documenting a high dynamic range and sensitivity. To control the specificity (above background) and on/off-kinetics of the reporter, we finally generated a doxycycline inducible short hairpin (sh)RNA mouse model targeting the endogenous prothrombin gene (Extended Data Fig. 2c). Upon doxycycline administration, this model (F2KD) reduces endogenous (pro)thrombin RNA, protein and activity down to 10-25% (Extended Data Fig. 2d). We next crossed this model with the reporter mice and determined luminescence with and without doxycycline administration. This analysis confirms the specificity of the Fluc signal in a prothrombin high abundant (liver) as well as in select low abundant tissues (kidney, spleen, Extended Data Fig. 2e,f). Further, the Fluc kinetics in the liver upon doxycycline addition and withdrawal corroborates the rapid on/off-kinetics of the reporter (Extended Data Fig. 2g), which is suited for real time imaging of dynamic expression alterations and resulting functional effects (Fig. 2b). Altogether this demonstrates the functionality of the multiplex imaging strategy to disentangle the expression and secretion dynamics of a prototypic liver-derived secretory protein in vivo. It also uncovers prothrombin expression in a variety of hitherto unknown extrahepatic locations.

Cell-based reporter systems are potent tools to elucidate disease mechanisms and to identify novel therapeutic vulnerabilities^51^. This becomes extremely efficient if such experimental set-ups can be carried out additively on the same genomic background. To test the in-principle applicability, we generated complementary syngeneic cell lines derived from our reporter animal. We used a modified extraction protocol^52^ to isolate primary hepatocytes from D-Insight and B6 control animals (Fig. 3, Extended Data Fig. 3a). After initial cultivation, we confirmed purity, iRFP and Fluc positivity of reporter hepatocytes using multispectral imaging flow cytometry (Fig. 3a, Extended Data Fig. 3b), ImageStream fluorescence microscopy (Fig. 3b, Extended Data Fig. 3c), luminometry and live cell imaging, where serial dilutions produced Fluc signals correlating with the number of cells (Fig. 3c). Recapitulating the findings obtained in vivo (Fig. 2a), we also detected Fluc and Rluc in hepatocyte cell culture lysates (Fig. 3d). Conversely, in the media only Rluc can be detected reflecting the secreted Rluc-tagged prothrombin, while Fluc remains cell-resident (Fig. 3d). Further, treating these cells with the protein transport inhibitor brefeldin A leads to a significant reduction of Rluc in the media (Fig. 3e), while, expectedly, the Rluc tagged-prothrombin accumulates in the cell. Together this confirms the functionality of this cell-based multimodal reporter imaging approach (Fig. 1a) enabling the unraveling of mechanisms governing the expression and secretion dynamics of this liver-derived secretory protein in high resolution and in a scalable format ex vivo. Additionally this multimodal approach provides multiple avenues for identification of novel cellular sources of prothrombin (e.g. flow cytometric analyses for the identification and functional characterization of prothrombin positive cells in the spleen and the lymphatic system (Fig. 2h) or hematopoietic (bone marrow) compartment (Extended Data Fig. 2b,3d,e).

**Figure 3.**
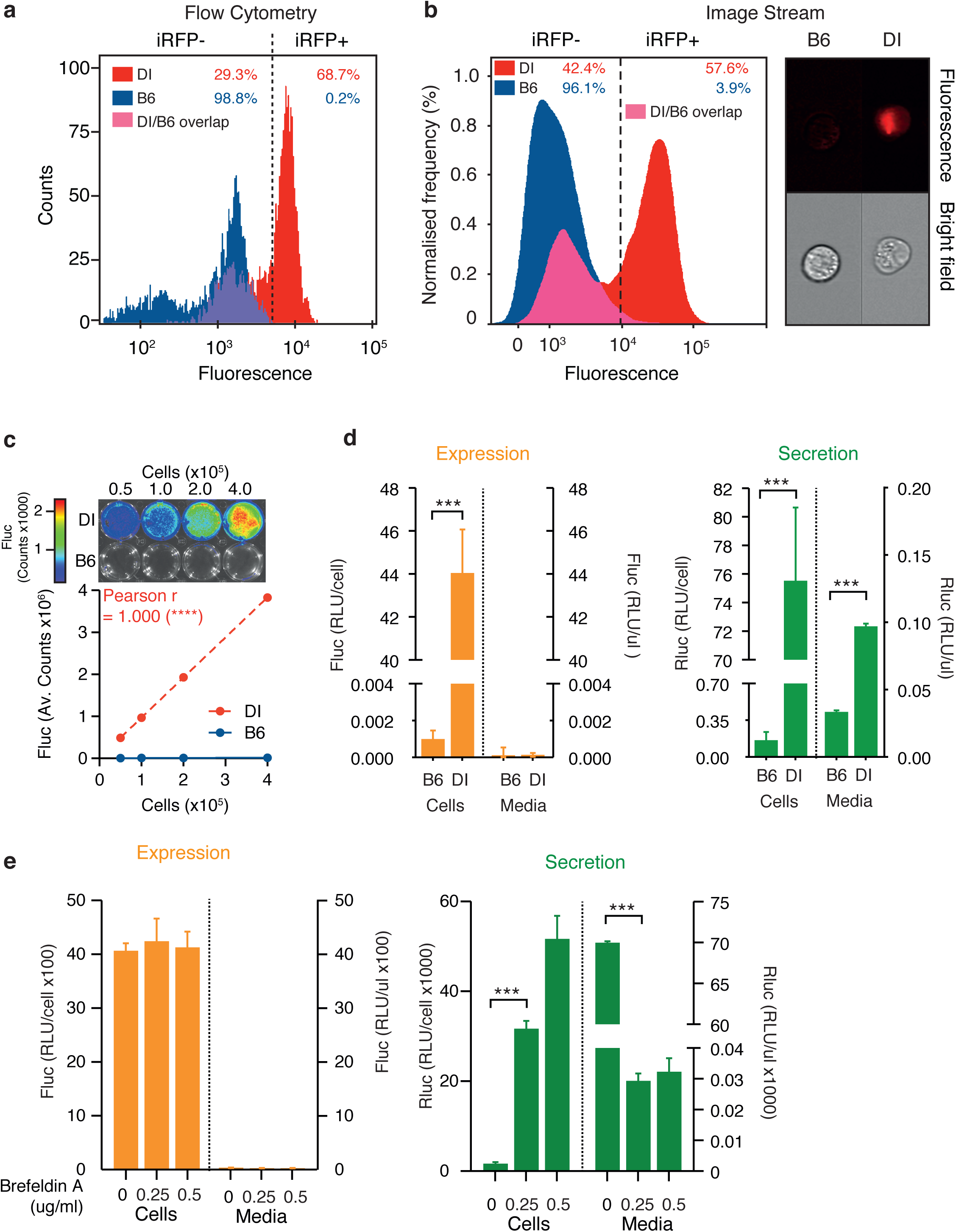
Complementary syngeneic primary D-Insight hepatocyte cell culture model for ex vivo exploration of the prothrombin expression and secretion dynamics. **a**, Flow cytometric (FACS) analysis and **b**, ImageStream microscopic flow cytometric analysis of primary hepatocytes isolated from D-Insight reporter and wildtype B6 control animals, documenting iRFP positivity of the reporter cell lines. **c**, Live cell imaging of primary D-Insight hepatocytes in culture shows a strong correlation between cell number and Fluc signal. **d**, Luciferase-luminometry confirms a specific Fluc and Rluc signal of primary D-Insight hepatocytes compared to wildtype B6 hepatocytes. It also reveals a specific Rluc signal in the cell culture supernatant (media) of the D-Insight hepatocytes, which reflects the Rluc-tagged and secreted prothrombin (green bars). **e**, Treatment of D-Insight hepatocytes with inhibitors of the secretory pathway (Brefeldin A) selectively reduces the Rluc signal in the cell culture supernatant (media), confirming the functionality and specificity of the genetic tagging strategy.

We next performed a proof-of-concept experiment to explore how the reporter mouse model performs in detecting potentially new locations of extrahepatic prothrombin expression with pathogenic relevance. Up to 20% of all cancer patients develop venous thromboembolism, reflecting a procoagulant disbalance of the hemostatic system with a significant incidence of fatal outcomes^53,54^. We therefore employed a methylcholanthrene (MCA) tumor induction model to assay whether prothrombin is potentially expressed and secreted by emerging tumors. In MCA challenged reporter animals, we first observed that the emerging tumors express prothrombin (Fig. 4a). We next transplanted these tumors into B6 mice and obtained Fluc signals that increase with time and volume of the transplanted tumor (Fig. 4b, Extended Data Fig. 4a,b). This suggests that the initial Fluc signals in the tumor unlikely originate from tumor-associated macrophages or lymphocytes frequently found in tumors^55^. To consolidate D-Insight-reporter and prothrombin expression, we analyzed tumor cell cultures obtained from these animals (Fig. 4c-f). Using multispectral imaging flow cytometry and luminometry, we confirm reporter iRFP positivity (Fig. 4c, Extended Data Fig. 4c) and Fluc expression (Fig. 4d). Using fibrosarcomas of the genetic background in which prothrombin expression can be depleted, we directly confirm endogenous prothrombin mRNA expression by F2 RNA-FISH (Fig. 4e,f). Surprisingly, probing further murine and human tumors we also find extrahepatic prothrombin expression in several entities (Extended Data Fig. 4d-f), suggesting that this finding may be relevant in tumor biology across species and entities^56^.

**Figure 4.**
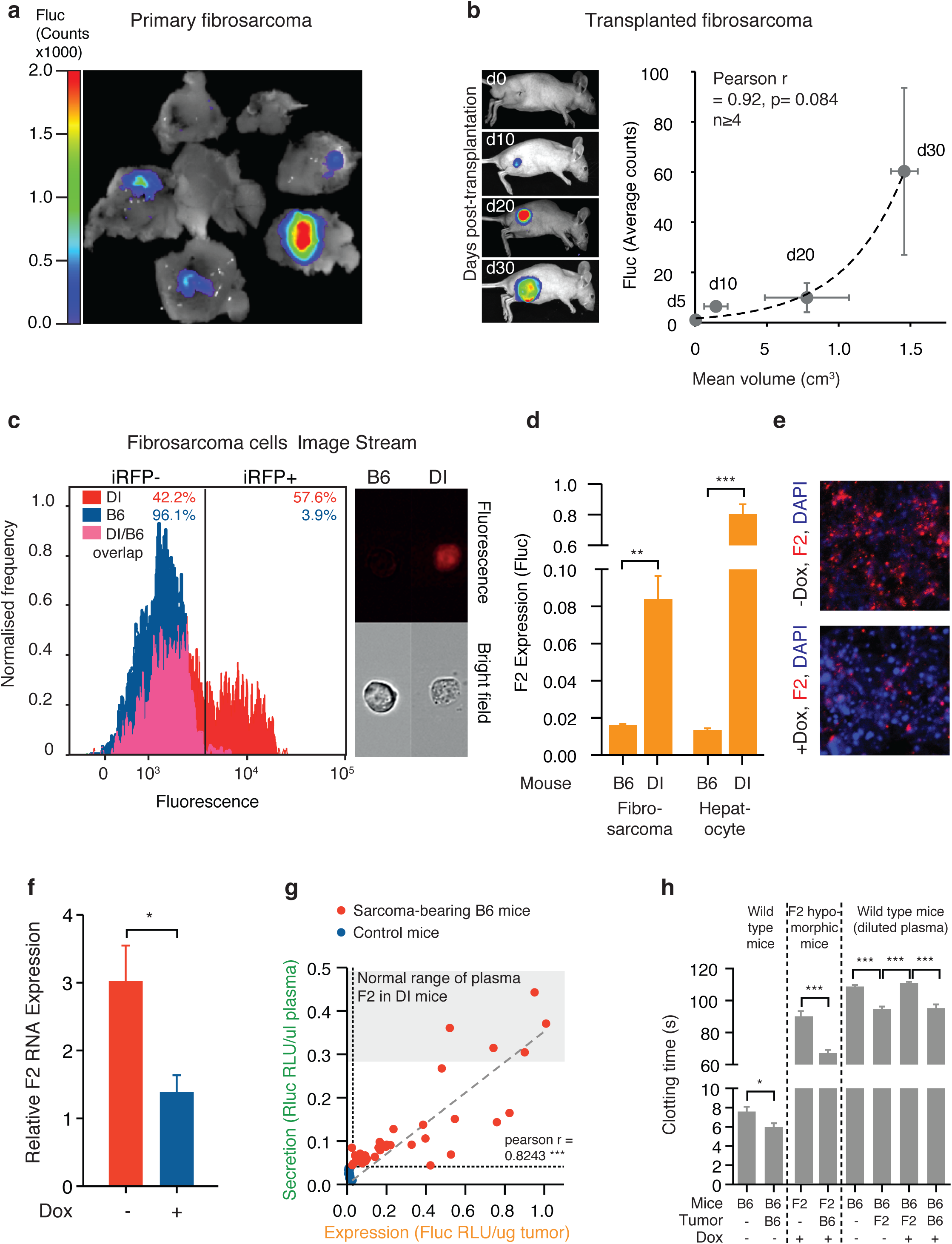
Identification of new locations of extrahepatic prothrombin expression with procoagulant function. **a**, MCA-induced fibrosarcomas in D-Insight mice, showing tumor-associated prothrombin expression (Fluc). **b**, Tumor-associated prothrombin expression is transplantable and increases with tumor volume. Non-invasive in vivo whole body bioluminescence imaging of Fluc of immunocompetent albino hairless (AH) mice after s.c. transplantation of D-Insight sarcomas at day 10, 20 and 30, respectively (y-axis depicts Fluc, the x-axis displays the tumor volume, n>3 animals per time point). Non-linear regression (Exponential growth equation) indicated by dashed line. **c**,**d**, ImageStream and luminometry of fibrosarcomas confirms iRFP positivity and Fluc expression for D-Insight fibrosarcomas. **e**,**f**, Tumor-intrinsic F2 expression confirmed by F2-RNA-FISH with and without doxycycline-mediated depletion of F2. **g**, Tumor-derived prothrombin is secreted into the blood circulation. The D-Insight reporter Fluc signal (x-axis) of sarcoma bearing mice (red) reflects the amount of prothrombin expressed in tumor cells, which positively correlates with the amount of tumor-derived prothrombin secreted into the circulation (y-axis, Rluc in plasma; signals obtained in muscle tissue and plasma, respectively, for B6 mice (blue) serve as non-specific background control). Notably, D-Insight sarcoma bearing B6 mice (red) express and secrete significant amounts of prothrombin, reaching Rluc plasma levels comparable to liver-derived prothrombin of D-Insight control mice (grey shaded corridor). **h**, Ectopically expressed prothrombin by the tumor is functionally active (assessed by KC4™ Coagulation Analyser; comparing blood plasma clotting time of F2-KD (F2), B6 mice (B6), F2-KD sarcoma bearing B6 mice and B6-sarcoma bearing F2-KD mice with (+) and without (-) administration of a doxycycline (Dox) rich diet for 3 weeks).

We next explored whether our model system provides further insights substantiating the functional dimension of this unexpected finding. We tested if tumor-derived prothrombin is secreted into the blood circulation. We transplanted D-Insight positive fibrosarcomas into B6 mice and determined Fluc and Rluc levels in the tumor and plasma, respectively. Surprisingly, we observed a strong correlation (Fig. 4g). Comparing these signals with Rluc in tumor-free D-Insight control animals reveals a substantial amount of tumor-derived Rluc that can be detected in the plasma, in concentrations comparable to tumor-free reporter animals (Fig. 4g). This suggests that tumor-derived prothrombin is quantitatively significant; it reaches the intravascular compartment to a similar extent as prothrombin constitutively synthesized in the liver.

The biogenesis of prothrombin involves post-translational gamma-carboxylation in the liver, which is required for full functional activity^57^. To determine the potential functionality of tumor-derived prothrombin, we first assessed whether all enzymes involved in the vitamin K cycle are expressed. We confirmed expression for the gamma carboxylase (GGXC), the NAD(P)H quinone hydrogenase 1 (NQOV1), the vitamin K epoxide reductase complex subunit 1 (VKORC1) and the vitamin K epoxide reductase complex subunit 1 like 1 (VKORC1L1) in the tumor at levels similar (or higher) to the liver (not shown). We also applied thrombin activity measurements and analyzed plasma samples obtained from either normal (B6) or prothrombin (F2) hypomorphic animals with and without implanted tumors that express prothrombin (Fig. 4f). To that end, we used wildtype B6 and F2 KD fibrosarcomas in which prothrombin expression can be selectively depleted (Fig.4e,f). We identified that tumor-derived prothrombin confers a procoagulant function in B6 mice (Fig. 4h, compare bars 1 and 2). Accordingly, we also observed that tumor-derived prothrombin compensates for low thrombin activity in prothrombin hypomorphic animals (Fig. 4h, bar 3 and 4), and that this compensation is lost, when the prothrombin expression in the tumor was silenced by the shRNA (Fig. 4h, bars 5-8). This suggests that tumor-derived prothrombin is functionally active. This may also perturb the well-balanced equilibrium between pro- and anticoagulatory activities in the plasma^58,59^ and could thereby contribute to the hypercoagulative state frequently observed in cancer patients, ultimately leading to detrimental consequences including thromboembolic death^53,54^.

Understanding tissue-specific gene regulation^60^ is key to tissue-tailored therapies in cancer and beyond^61^. Based on our findings, a therapeutic strategy targeting the expression of such a secretory protein in a tissue-selective manner would be desirable. We therefore finally tested the applicability of this approach and conducted a small-scale chemical screening employing a culture of primary hepatocytes, fibrosarcoma tumor cells and fibroblasts (serving as controls) obtained from the D-Insight reporter animal (Fig. 3,4). Using in-plate luminometry in a 96-well format in our screening, we surprisingly identified a DNA methyltransferase inhibitor (Decitabine), which shows a long-lasting and highly selective modulation of prothrombin gene expression in a tissue dependent-manner affecting fibrosarcomas, while leaving the expression in the liver and fibroblasts unaltered (Fig. 5, heatmap and Extended Data Fig. 5). This points to tissue-selective regulatory mechanisms governing the expression of this secretory protein with a multitude of actions in various systems and disease mechanisms, ranging from atherosclerosis to tumor biology^35,36^. It also illustrates the versatility of this model to dissect disease mechanisms and to uncover novel therapeutic vulnerabilities (Fig. 5).

**Figure 5.**
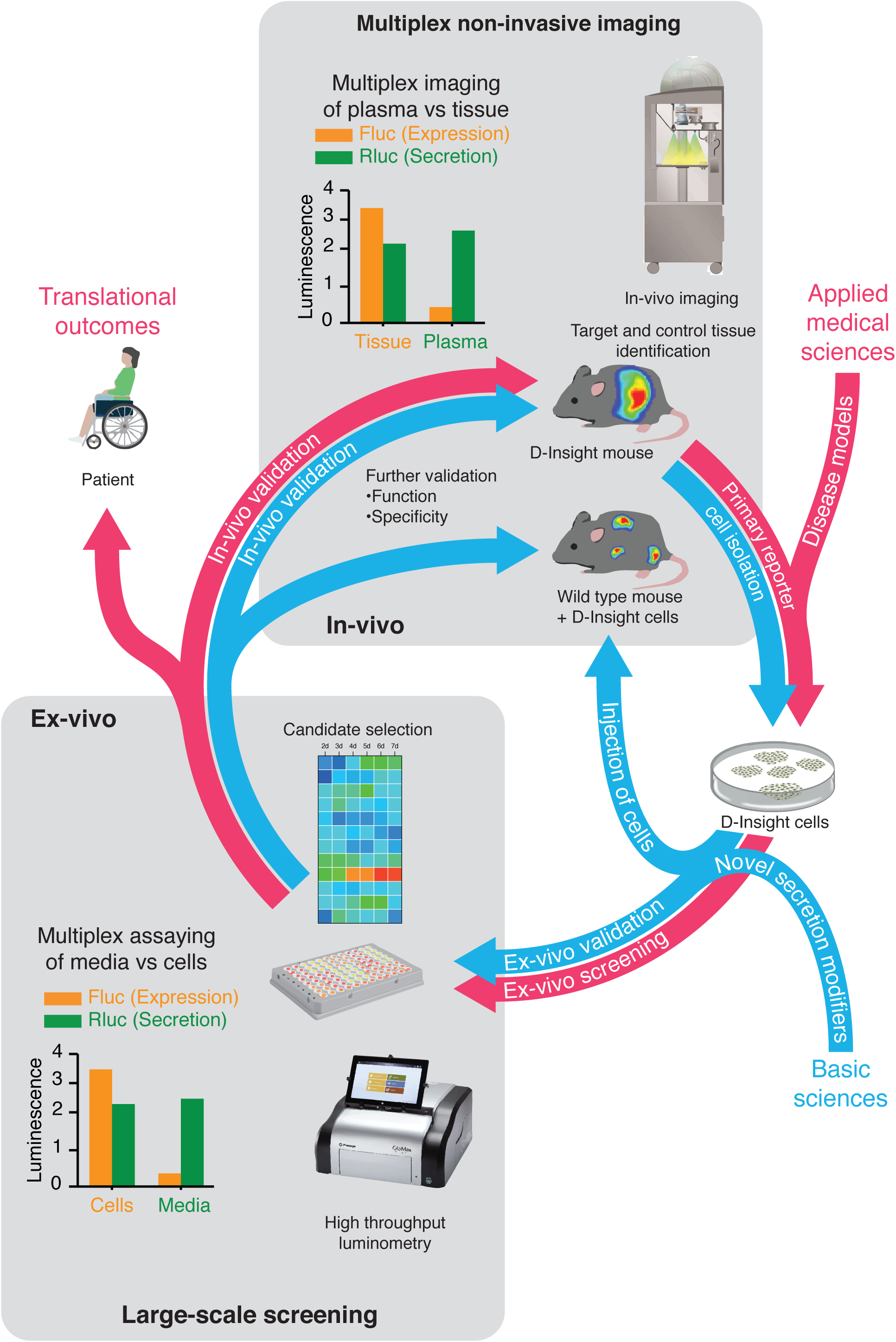
An integrated genetic reporter system for basic and applied medical research of secretory proteins. Exemplary representation of a complementary reporter model (screening) approach based on multiplex non-invasive in vivo imaging in combination with the isolation and cultivation of D-Insight-reporter positive target cells (e.g. hepatocytes). Reverse and forward pharmacology approach unified in syngeneic reporter set-up. Here cells endogenously expressing and secreting prothrombin, as well as cells induced in disease models, are identified by non-invasive imaging, isolated and cultured. These can be screened ex-vivo through high-throughput luminometry, and additionally in-vivo, through injection into wild-type mice and non-invasive imaging. Findings made in these systems can finally be validated in-vivo using the original D-Insight model. The heat map in the center exemplifies a tissue selective targeting, obtained from a pilot proof-of-concept study (shown in Extended Data Fig. 5).

## Discussion

The liver is the prime organ producing secretory proteins. While protein targeting is achieved by a well-characterized cargo system^3,4^, new mechanisms of unconventional protein secretion are being discovered^62,63^. Owing to the importance of secretory proteins in a variety of diseases^9-14^, understanding regulatory mechanisms on a systems and cellular level may help elucidating underlying biology and pathophysiological processes, including the identification of novel therapeutic avenues^23^.

Here we introduce a conditional, multiplex imaging reporter mouse model to disentangle the expression and secretion dynamics of a prototypic liver-derived secretory protein in vivo. Our findings illustrate the versatility of this reporter model for detecting and illuminating processes of this secretory protein in the context of development and various disease mechanisms. They also document the autonomous expression of a secretory protein that becomes harmful if aberrantly expressed. Additionally, our pilot screening demonstrates that this reporter tool set is suited to uncover tissue-specific regulatory mechanisms that may allow tailored therapeutic targeting. Given the broad expression of prothrombin in various tissues, the reporter model introduced here will likely find wide application to understand and dissect the role of prothrombin in numerous extravascular pathophysiologies^36,64^.

Biologically, our proof-of-concept observations made here reveal a novel mechanism as to how cancer contributes to a hypercoagulant phenotype. Overexpression of prothrombin perturbs the well-balanced equilibrium between pro- and anticoagulatory activities in the plasma. While the hyperexpression observed in tumors is low level, the cumulative secretion appears to quantitatively exceed the procoagulant effects caused by the well-established thrombophilic prothrombin polymorphism F2 20210 G>A ^65,66^. Even small tumor lesions can lead to the hypercoagulative state frequently observed in cancer patients, ultimately leading to detrimental consequences including thromboembolic death^53,54^. In view of the evolution of direct oral anticoagulants (DOACs), charting the spatiotemporal dynamics of prothrombin in pathophysiologies may also reveal hitherto unrecognized advantages of selective therapeutic targeting of the hemostatic system^67^.

Beyond discovering new functions of the hemostatic system with spatiotemporal resolution, this reporter model is multifunctional. It allows exploring fundamental biological principles of gene regulation and protein targeting, and to identify rheostats involved in the cross-talk between gene expression and availability of a secretory protein. Complementary syngeneic primary cell culture models also enable the deconvolution of regulatory mechanisms in high resolution, and in a scalable high throughput format ex vivo. It thereby represents a valuable resource for uncovering disease-eliciting cues and novel therapeutic vulnerabilities, including their validation by non-invasive imaging in vivo, and vice versa (Fig. 5). Ultimately this model is an instructive example for a successful multimodal imaging strategy, which can be simply adapted to numerous other secretory proteins.

## Acknowledgements

The authors would like to express their gratitude to current and former members of the Danckwardt lab. We are indebted to Enrico Di Cera for sharing his thoughts on the initial concept of the prothrombin-fusion construct, and Anca Dragulescu-Andrasi and Andreas Loeing for their supportive information on the modified Rluc proteins. Kathrin Mühlbach for her supportive discussion on the targeting strategy, Friederike Häuser on protocols of hepatocyte isolation, and Kristian Schütze, Antje Canisius and Kevin Friedemann for technical support.

This work is supported by grants of the DFG (DA 1189/2-1), the GRK 1591, the DFG Priority Program SPP 1935 (Deciphering the mRNP code: RNA bound Determinants of Post-transcriptional Gene Regulation), by the Federal Ministry of Education and Research (BMBF01EO1003), by the Hella Bühler Award for Cancer Research, and by the German Society of Clinical and Laboratory Medicine (DGKL).

## Contributions

JN, ST, SK, EK, LKS, LH, LE, DAS performed experiments. JN and ST analyzed and compiled data. HC interpreted data and discussed results. SD conceived and supervised the study, performed experiments and wrote the paper with contributions from all coauthors.

## Material and Methods

### Generation of the D-Insight mouse line

The targeting strategy aimed at generating a constitutive Knock-In (KI) of Rluc (Renilla Luciferase)-P2A-Flag-iRFP (near-infrared fluorescent protein)-P2A-DTR (diphtheria toxin receptor)-P2A-Fluc2CP (Firefly Luciferase) at the F2 gene (NCBI transcript NM_010168_2). To that end, the following cassette has been inserted between the last amino acid and the translation termination codon in exon 14 of the F2 gene (from 5’ to 3’, Supplementary Figure S1):

- an in frame loxP site (nucleotide sequence: ataacttcgtatagcatacattatacgaagttatgc)
- a GGGSGGGGSGGGGSGGGGS-linker peptide (nucleotide sequence: ggcggcggcagcggcggcggcggcagcggcggcggcggcagcggcggcggcggcagc)
- the RLuc 8.6 open reading frame (ORF)
- the “self-cleaving” peptide P2A (nucleotide sequence: gctactaacttcagcctgctgaagcaggctggagacgtggaggagaaccctggacct)
- a Flag tag sequence (nucleotide sequence: gactacaaggacgacgatgacaag)
- a GGGSGGGS-linker peptide (nucleotide sequence: ggcggcggcagcggcggcggcagc)
- the iRFP ORF
- a GGGSGGGS-linker peptide (nucleotide sequence: ggcggcggcagcggcggcggcagc)
- the “self-cleaving” peptide P2A (nucleotide sequence, see above)
- the DTR ORF
- an in frame loxP site (nucleotide sequence: ataacttcgtatagcatacattatacgaagttatat)
- the “self-cleaving” peptide P2A (nucleotide sequence, see above)
- the Fluc2CP ORF.

Prior to cloning of the targeting vector, all inserted sequences have been tested and modified for optical codon usage (in mice). All used linker peptides were tested for potential interference with secondary and tertiary structures of F2, RLuc 8.6, iRFP, DTR and Fluc2CP. The positive selection marker (Puromycin resistance - PuroR) has been flanked by FRT sites and inserted into intron 12. The targeting vector has been generated using BAC clones from the C57BL/6J RPCIB-731 BAC library and has been transfected into the C57BL/6N Tac ES cell line. Homologous recombinant clones have been isolated using positive (PuroR) and negative (Thymidine kinase - Tk) selection. Constitutive KI allele 1 was obtained after Flp-mediated removal of the selection marker. This allele expressed a chimeric transcript encoding F2 protein fused to the loxP-Rluc ORF, the P2A sequence, the Flag tag sequence fused to the iRFP ORF, the P2A sequence, the DTR-loxP fusion protein, the P2A sequence and the Fluc2CP ORF. The expected co-translational „cleavage” at the P2A sequences results in a 1:1:1:1 stochiometric co-expression of the F2-loxP-Rluc, Flag-iRFP and DTR-loxP fusion proteins and the Fluc2CP protein under the control of the endogenous F2 promoter.

Constitutive KI allele 2 was obtained after Cre-mediated removal of the Rluc, the Flag-iRFP and the DTR ORFs. This allele should express a chimeric transcript encoding F2 protein fused to the loxP and the P2A sequences and the Fluc2CP ORF. The expected co-translational cleavage at the P2A sequences should result in co-expression of the F2-loxP fusion protein and the Fluc2CP protein under the control of the endogenous F2 promoter. The remaining recombination site is located in a non-conserved region of the genome.

For conditional imaging heterozygous D-Insight animals were paired with Albumin-CRE (Alb-Cre) animals (B6.Cg-Speer6-ps1Tg(Alb-cre)21Mgn/J)^68^.

### Inducible F2-RNAi mouse model

For reversible depletion of F2, an inducible knock-down allele of the F2 gene was generated via targeted transgenesis of a doxycycline-inducible shRNA cassette into the ROSA26 locus (Gt(ROSA)26Sor)^69^. Briefly, to that end a recombination-mediated cassette exchange vector harboring an inducible H1 promoter (H1tetO)-driven shRNA cassette along with a genetic element for the constitutive expression of the codon-optimized tetracycline repressor protein (iTetR), and a neomycin resistance cassette was transfected into C57BL/6 ES cell line equipped with RMCE docking sites in the ROSA26 locus. Recombinant clones were isolated using neomycin resistance selection and positive clones harbouring six different shRNAs targeting F2 were pretested for knockdown potency in ES cells (by qPCR analysis). The clone with highest knock-down efficiency was used for the generation of the mouse line. All animal experiments were approved by local authorities, and animals’ care was in accordance with institutional guidelines.

### Isolation of primary hepatocytes and primary hepatocyte culture

Mice were anesthetized with Ketamine (50 μg/ml, Inresa Arzneimittel GmbH) and the abdomen was opened. Subsequently the portal vein was cannulated to perfuse the liver with 100 ml of prewarmed (37°C) primary hepatocyte isolation (PHI) buffer I (137 mM NaCl, 52 mM KCl, 10 mM HEPES, pH7.4). Upon successful flushing, 50 ml of PHI-buffer II (58 mM NaCl, 52 mM KCl, 1.6 mM CaCl_2_, 10mM HEPES, 0.05% collagenase, pH7.6) was used for digestion of the liver. Afterwards, the perfused liver was explanted and stored in PHI-buffer III (137 mM NaCl, 52 mM KCl, 1.6 mM CaCl_2_, 10 mM HEPES, pH7.6). The liver was then mechanically dissociated under sterile conditions and dissociated through a 40 μM cell strainer (Corning) and centrifuged (90 g, 3 minutes without break). Next the supernatant was discarded and the liver cells resuspended in 15 ml prewarmed PBS followed by a second round of centrifugation (300 g, 3 minutes without break). The cells were resuspended in William’s E medium (Biochrom). Viability was measured by the trypan blue exclusion test. Hepatocytes were then seeded on gelatin-coated culture plates (0.2% gelatin) for 1 hour at 37°C in William’s E medium containing 5 % FBS, 1 μM Dexamethasone in DMSO, 100 units/ml Penicillin, 100 μg/ml Streptomycin, 4 μg/ml Human Recombinant Insulin, 2 mM GlutaMAX, 15 mM HEPES at 37 °C in/5%CO_2_. The culture medium was changed every 2-3 days. D-Insight positivity of primary hepatocytes was confirmed by luminometric analysis (see below), by FACS based on iRFP fluorescence in a BD FACS Canto (BD Bioscience) and by an Amnis ImageStream Mk II flow cytometer (Luminex) and analyzed using the IDEAS software (Luminex).

### Generation and assessment of MCA-induced tumors

Methylcholanthrene (MCA) induced sarcomas were generated through 0.1mg MCA suspension (Sigma #213942) in 30μL corn oil (Sigma #C8267) intramuscularly injected into the left gluteal muscle of D-Insight animals (8-12 weeks old).

### Generation of tumor cell lines

Tumors were extracted before they reached diameters >2cm. The tumors were washed in PBS and cut into several fragments and transferred into a 6-Well-Plate containing digestion medium (RPMI, 20 % FBS, 100 units/ml Penicillin, 100 μg/ml Streptomycin, 8 mg/ml Collagenase) and incubated for 24 hours at 37 °C in 5%CO_2_.umor fragments were mechanically dissociated by pipetting up and down, and then filtered through 40 μm cell strainers. The suspension was centrifuged at 1000 x g for 5 minutes and the obtained cell pellet was resuspended in DMEM (Biochrom). Viability was measured by the trypan blue exclusion assay. Fibrosarcoma cells were seeded on culture plates in DMEM containing 20 % FBS, 3.7 g/l NaHCO_3_ 4.5 g/lD-Glucose at 37 °C in 5%CO_2_. The medium was changed every 48 h.

### Generation of primary fibroblast cultures

Mice ears were washed in PBS and cut into several fragments. The fragments were transferred into a 6-well-plate containing digestion medium (RPMI, 20 % FBS, 100 units/ml Penicillin, 100 μg/ml Streptomycin, 8 mg/ml Collagenase) and incubated for 24 hours. Ear fragments were mechanically dissociated by pipetting up and down and filtered through 40 μm cell strainers. The suspension was centrifuged at 1000 x g for 5 minutes and the obtained cell pellet was resuspended in DMEM. Viability was measured by the trypan blue exclusion assay. Fibroblasts were seeded on culture plates in DMEM containing 20% FBS, 3.7 g/l NaHCO_3_, 4.5 g/l D-Glucose at 37 °C in 5%CO_2_. The medium was changed every 48 h.

### In vivo Imaging

Non-invasive in vivo imaging of D-Insight mice was performed under Ke/Xy narcosis (intraperitoneal (i.p.) injection of ketamine (Ke, 80-120 mg/kg body weight) / xylazine (Xy, 5-10 mg/kg body weight) cocktail) using an IVIS Spectrum In Vivo Imaging System (Perkin Elmer, Rodgau, Germany) with images acquired using the Living Image® software package (Perkin Elmer).

The dorsal or ventral images of mouse bodies, ventral images of mice in operation, ex vivo imaging of the same mice (after euthanasia by overdose application of Ke/Xy narcosis) or postmortem imaging of certain organs were performed 30 min after i.p. injection of 100 μl ready-to-use coelenterazine (RediJet Coelenterazine H Bioluminescent Substrate, PerkinElmer, Rodgau, Germany) or 10 min after i.p. injection of 100 μl ready-to-use luciferin (RediJet D-Luciferin Bioluminescent Substrate, PerkinElmer, Rodgau, Germany) respectively. The pseudo color luminescent images (blue, green, yellow, and red from least to most intense) were overlaid on the grayscale photographic images. Quantification of luminescence was determined by calculating the Average Counts as Total Counts/Number of pixels for a defined region of interest.

### Luminometry Assays

100000 cells or 200 mg tissue were lysed in 100-500 μl Passive Lysis Buffer (Promega) for 15 min at room temperature on an orbital shaker. 50 μl of the lysate, plasma or media samples, were assayed in white 96 well plates in triplicate by the addition of 50 μl Bright-Glo™ Firefly luciferase assay system (Promega) or freshly prepared 40 ng/ml Coelenterazine in Renilla-Glo buffer (Promega). Alternatively 20 μl samples were assayed using a Dual-Glo Luciferase Assay System using manufacturers protocol (Promega). Luminescence (RLU) was measured immediately using a GloMax® Discover Microplate Reader (Promega).

### Protein Transport Inhibitor Treatment

200000 cells were seed in Hepatocytes-Williams E medium (Gibco, 10043282) containing 10% FCS, 100 units/ml Penicillin and 100 μg/ml Streptomycin and incubated at 37 °C in 5% CO2. After 3 h, 0-0.5 ug/ml Brefeldin A was added to the medium and cells were further incubated for 2 days. Luminometry assays were then performed as described above.

### Image Stream

For real time imaging of iRFP signal in DI hepatocytes and fibrosarcoma cells image stream was used. Cells were trypsinized and diluted in FACS buffer (PBS (1x), Bovine serum albumin (BSA, 0.5%), fetal bovine serum (FBS, 0.1%), Sodium azide (NaN3, 0.1%)) to a final concentration of 1 million cells per 500 μL. B6 hepatocytes and fibrosarcoma cells were used as controls. 1 μL of DAPI (1:10000 diluted in PBS) was added to 20 μL cell suspension before loading samples on the ImageStreamX MkII imaging cytometer. Single cell population and live cells were gated by considering the signal generated in DAPI 405 channel. For activation of iRFP signal in fibrosarcoma cells, 3-5 μL of biliverdin hydrochloride (chromophore for iRFP signal activation, final concentration 0.002%, Sigma; 30891) added to cells and incubated for 10 mins at 37 °C before imaging. As liver cells are known to produce sufficient amounts of bilirubin no additional biliverdin was used for iRFP activation in hepatocytes population. Living cells were sorted for iRFP positive signal and were quantified by gating the cells based on signal from 702/86 channel. The INSPIRE software was used to analyze the data and graph prism was used to plot the graphs.

### LPS-treatment

Mice were injected intra-peritoneal with 1.0 to 1.5 mg/kg LPS (Sigma Aldrich). After 3 to 48 h mice were sacrificed by isoflurane inhalation and liver and plasma samples obtained. Samples were assayed by luminometry as described above.

### F2 qRT-PCR

Total RNA recovered from cells using peqGOLD TriFast reagent (Peqlab) and cDNA produced using the RevertAid Reverse Transcriptase kit (ThermoFisher Scientific) as per manufacturer’s instructions. qRT-PCR was performed using the Blue S’Green qPCR Kit (Biozym) with primers for F2 (GGTGAACCTGCCCATTGTAGA, TCCTCGCTTGGTGTCATTCA) and the housekeeper gene GAPDH (AGGTCGGTGTGAACGGATTTG, TGTAGACCATGTAGTTGAGGTCA) as per manufacturer’s instructions. Cycling (95°C for 2 min; 95°C for 30 sec; 60°C for 30 sec; 72°C for 30 sec and followed by a melting curve) was performed in a CFX Real-Time Detection System (Bio-Rad) and relative expression calculated using the CFX Manager Software (Bio-Rad).

### F2 RNA in situ hybridization (FISH)

Cryosections were generated from murine organs and tissues. Briefly, mice were sacrificed by isoflurane inhalation and organs were washed in PBS before placing in Tissue-Tek^®^ Cryomolds (25×20×5mm or 15×10×5mm; Sakura). The specimens were immediately covered with Tissue-Tek^®^ O.C.T. compound, put on dry ice until frozen and stored at −80°C until sectioning. Cryosections (12 μm) were produced using a cryostat, mounted onto Superfrost^®^plus slides (Thermo Fisher Scientific) and fixed overnight at 4°C in 4% formaldehyde (FA).

The Thermo Fisher Scientific ViewRNA ISH Tissue 1-Plex Assay was used to visualize the F2 mRNA expression in cryosections. Briefly, slides were washed twice in PBS and dehydrated in increasing EtOH concentrations (50%, 70%, 100%; 10 min each). After drying, a hydrophobic barrier was drawn around the specimen using the ImmEdge pen (Vector). Slides were boiled (85-95°C) for 30-45s in pretreatment solution and washed twice with PBS, followed by incubation in 0.2M HCL (RT for 10 min). After washing with PBS, slides were incubated (10 min, RT) with Protease solution (diluted 1/100 in prewarmed PBS), followed by rinsing in PBS and 3 minutes incubation in 4% FA. Thereafter, all following hybridization and amplification steps were carried out at 40°C in a humidified chamber.

After washing twice with PBS and once with ddH_2_O, the hybridization of the probe set was carried out by incubation with the probe set working solution (probe set diluted 1/30 with prewarmed Probe set diluent) for 2.5-3h at 40°C. After hybridization, slides were washed 3x in washing buffer for 2 min at room temperature and under vigorous shaking. The slides were then incubated for 40 min in PreAmp solution diluted 1/100 in Amplifier Diluent, followed by a second washing step as indicated above. The slides were then incubated for 40 min in Amp solution diluted 1/100 in Amplifier Diluent. After a third washing step, the slides were incubated in Label Probe-AP diluted 1/500 in Label Probe Diluent for 40 min, followed by washing 3x in washing buffer for 3 min each with continuous shaking. AP-Enhancer solution was added to the samples and incubated for 5 min at room temperature. The slides were then incubated with freshly prepared detection substrate, consisting of half a tablet of Fast Red Substrate resuspended in 2.5 ml Naphthol buffer, for 30-40 min at 40°C in the humidified chamber. Slides were then washed 2x in PBS and kept in 4% FA for 5 min. After washing in ddH_2_O, counterstaining in haematoxylin for 10-30 sec and destaining in ddH_2_O, slides were dried 20 min at room temperature in dark environment, mounted with Fluoromount-G with DAPI (Invitrogen), covered with coverslips and dried again for 15 min. Nail polish was applied to seal the edges and slides were stored at 4°C covered with aluminum foil.

### F2 activity, APTT measurements, Clinical Chemistry

F2 activity of tumor-bearing and tumor-free animals was measured with ACL Top instruments and a KC4™ Coagulation Analyser. Analyses of transaminases (GOT/GPT) have been carried out on an Abbott Alinity C analyzer.

### Diphtheria-Toxin treatment

The dose of Diphtheria toxin (DTX) was titrated in the range 0-4 ng/g body weight per animal with 2 ng/g being defined as optimal and used for the shown experiment (injected i.p.).

### Epigenetic modulator treatment of D-Insight fibrosarcoma cells

100000 D-Insight fibrosarcoma cells were seeded in 6 Well plates in DMEM containing 10% FBS, 3.7 g/l NaHCO3, 4.5 g/l D-Glucose and incubated at 37 °C in 5% CO2 for 24 hours. Media was then changed to include an epigenetic modulator (either 10 nM (+)JQ1, 10 μM Azacitidine, 10 μM Decitabine, 10 μM Lomeguatrib, 10 nM Panobinostat, 10 nM Quisinostat, 10 μM RG108 or 10 μM Zebularine in DMSO) and incubated for 2 - 7 days and luminometry performed as described above.

### Statistical Analysis

All statistical analyses and production of graphs were performed using Prism 9 (GraphPad Software).

---

**Extended Data Figure 1.**
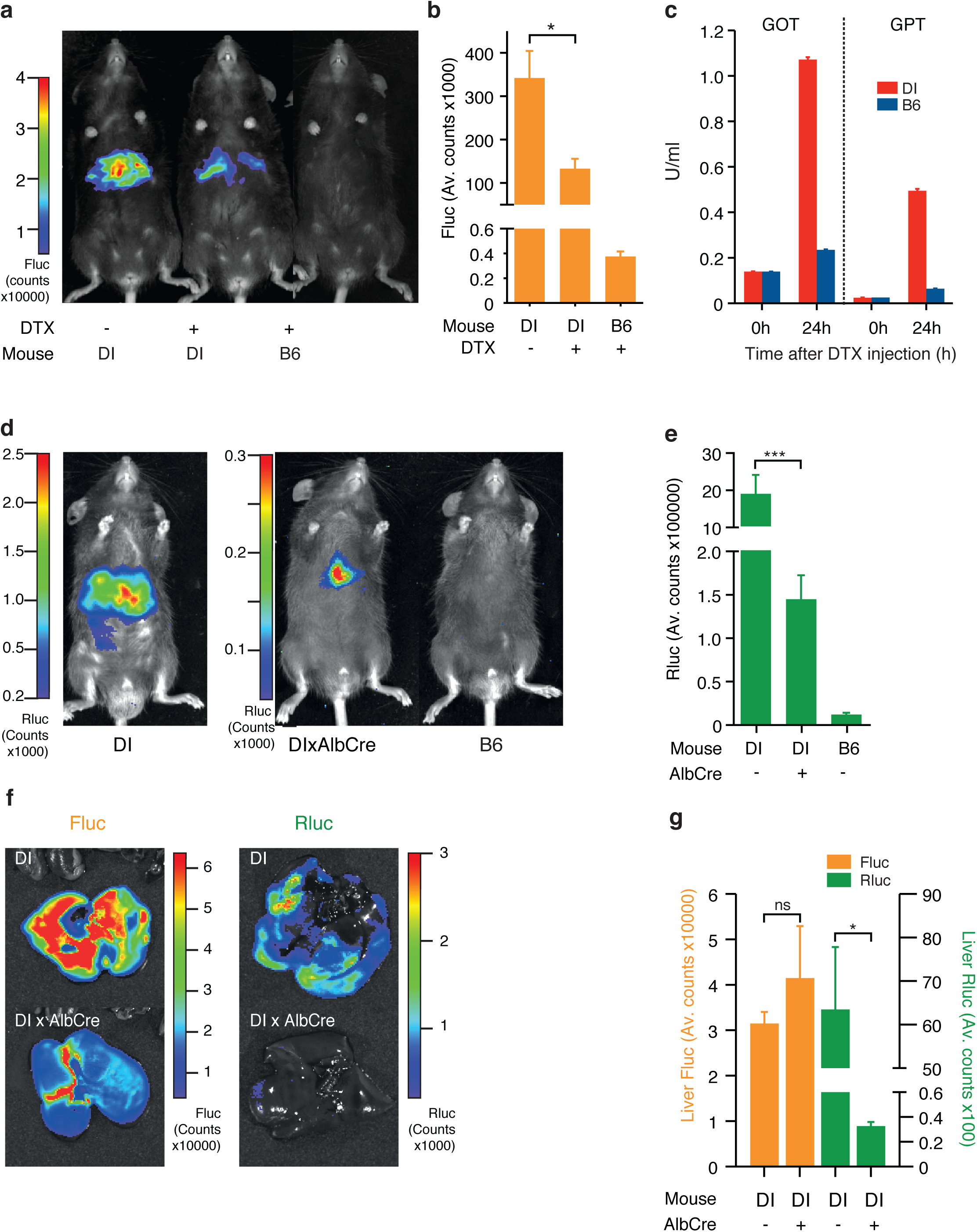
Functional customization of the D-Insight model through Diphtheria toxin receptor mediated targeting and Cre-mediated reporter excision. **a**-**c** Validation of Diphtheria toxin (DTX) mediated targeting of reporter expressing cells (DI +/-DTX), showing reduction of liver Fluc signal **b**, and a corresponding release of liver derived transaminases (glutamic oxaloacetic transaminase (GOT), glutamic pyruvate transaminase (GPT)) 24 h after DTX injection in comparison to a wildtype B6 animal **c. d**-**g** Validation of Cre-mediated excision of the Rluc-DTR reporter cassette with D-Insight animals after pairing with a liver specific Cre-line (AlbCre). This results in a significant reduction of liver-derived Rluc bioluminescence (**e, f, g**), confirming functionality of the conditional multiplex reporter labeling strategy (Fig. 1a).

**Extended Data Figure 2.**
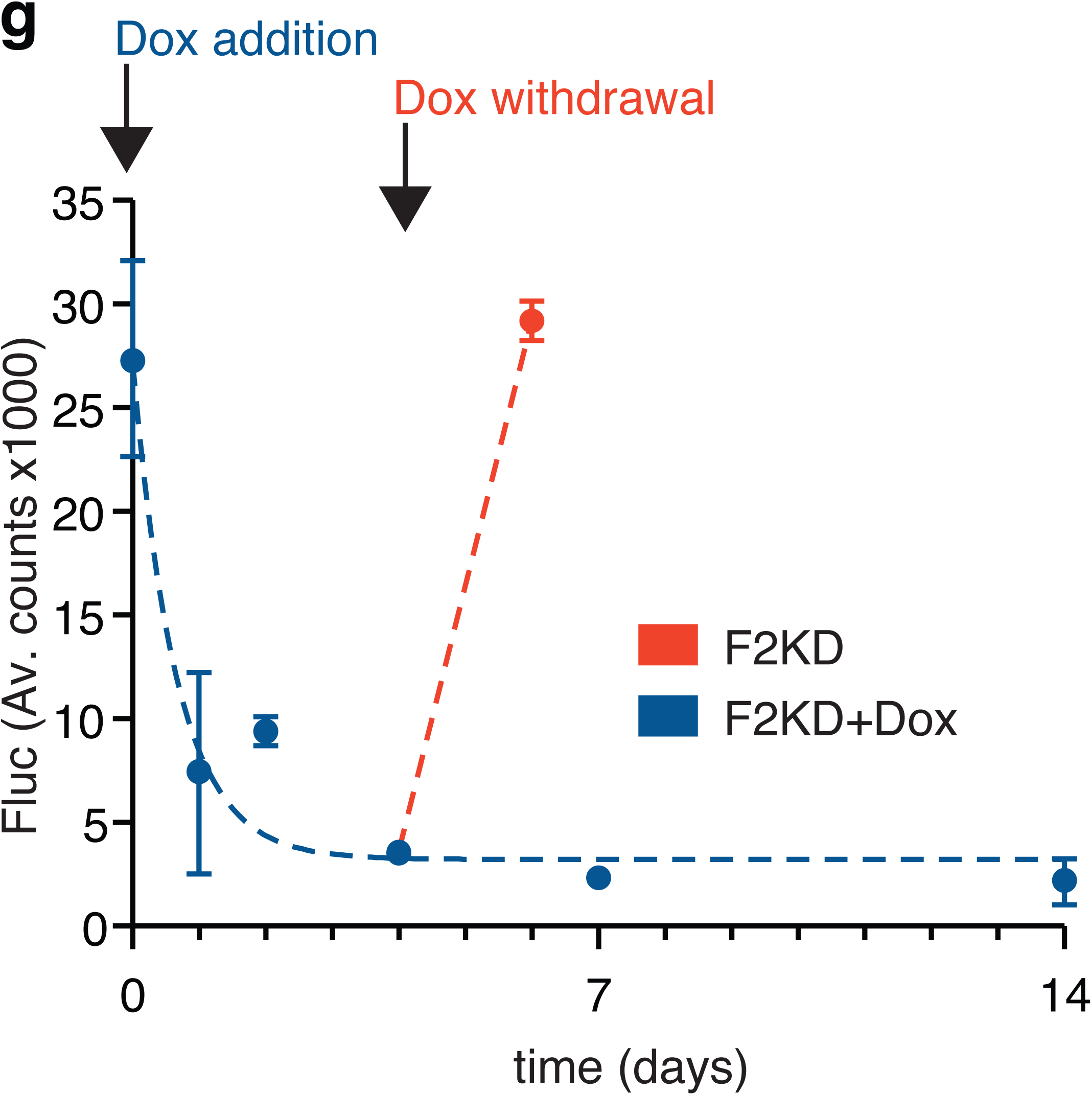
Performance measures of the D-Insight reporter to track (dynamic changes of) prothrombin gene expression. **a**, Functionality of Rluc tagged prothrombin compared to untagged prothrombin (B6), confirming (near) complete thrombin activity (clotting time) in D-Insight animals. **b**, Comparison of tissue lysate luminometry (Fluc, y-axis) with a prothrombin specific qRT-PCR (x-axis), confirming a high sensitivity, accuracy and dynamic range of the D-Insight reporter model. **c**, Schematic of the doxycycline inducible shRNA mouse model (F2KD) to silence prothrombin. **d**, Doxycycline-mediated induction of shRNA reduces prothrombin mRNA and protein in the liver, and prolongs the time to clot (i.e. reduces the thrombin activity). **e**,**f** Doxycycline-mediated induction of shRNA reduces Fluc signals in double-heterozygous reporter animals (F2KDxD-Insight) in prothrombin high- (liver) and prothrombin low- (spleen, kidney) abundant tissues, corroborating the specificity of the recorded Fluc reporter signals. **g**, Reporter on/off kinetics (Fluc) in living double-heterozygous reporter animals (F2KDxD-Insight) after addition and termination of doxycycline supplementation. Non-linear regression (One phase decay) indicated by dashed line.

**Extended Data Figure 3.**
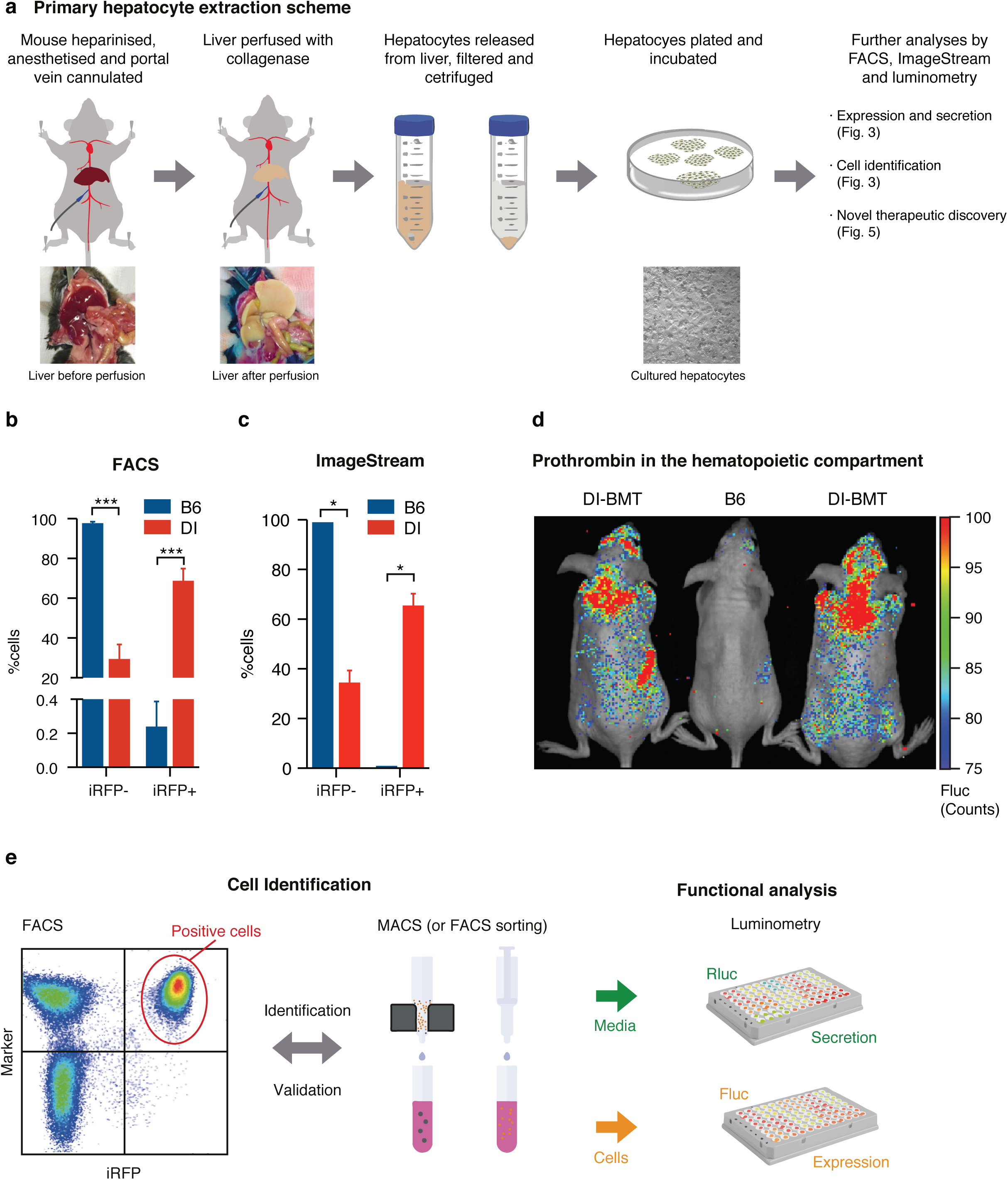
Generation of primary reporter cell cultures for large-scale identification of expression and secretion modifiers. **a**, Schematic for primary hepatocyte cell isolation. **b-c**, FACS and ImageStream based confirmation of iRFP positivity of viable hepatocytes obtained from D-Insight reporter and B6 control animals. **d**, Hematopoietic prothrombin in B6 animals transplanted with D-Insight bone marrow (DI-BMT) as compared to a B6 control, corroborating Fluc positivity in the bone marrow (Extended Data Fig. 2b) **e**, Schematic of the cell identification (and validation) of prothrombin expressing cells based on iRFP in flow cytometric analyses (left) and enrichment analysis (MACS or FACS sorting) followed by luminometry. For functional analyses, cultured cells (right) serve as reporter models for (large scale) identification of expression and secretion modifiers (Fig. 3, 5, and Extended Data Fig. 5)

**Extended Data Figure 4.**
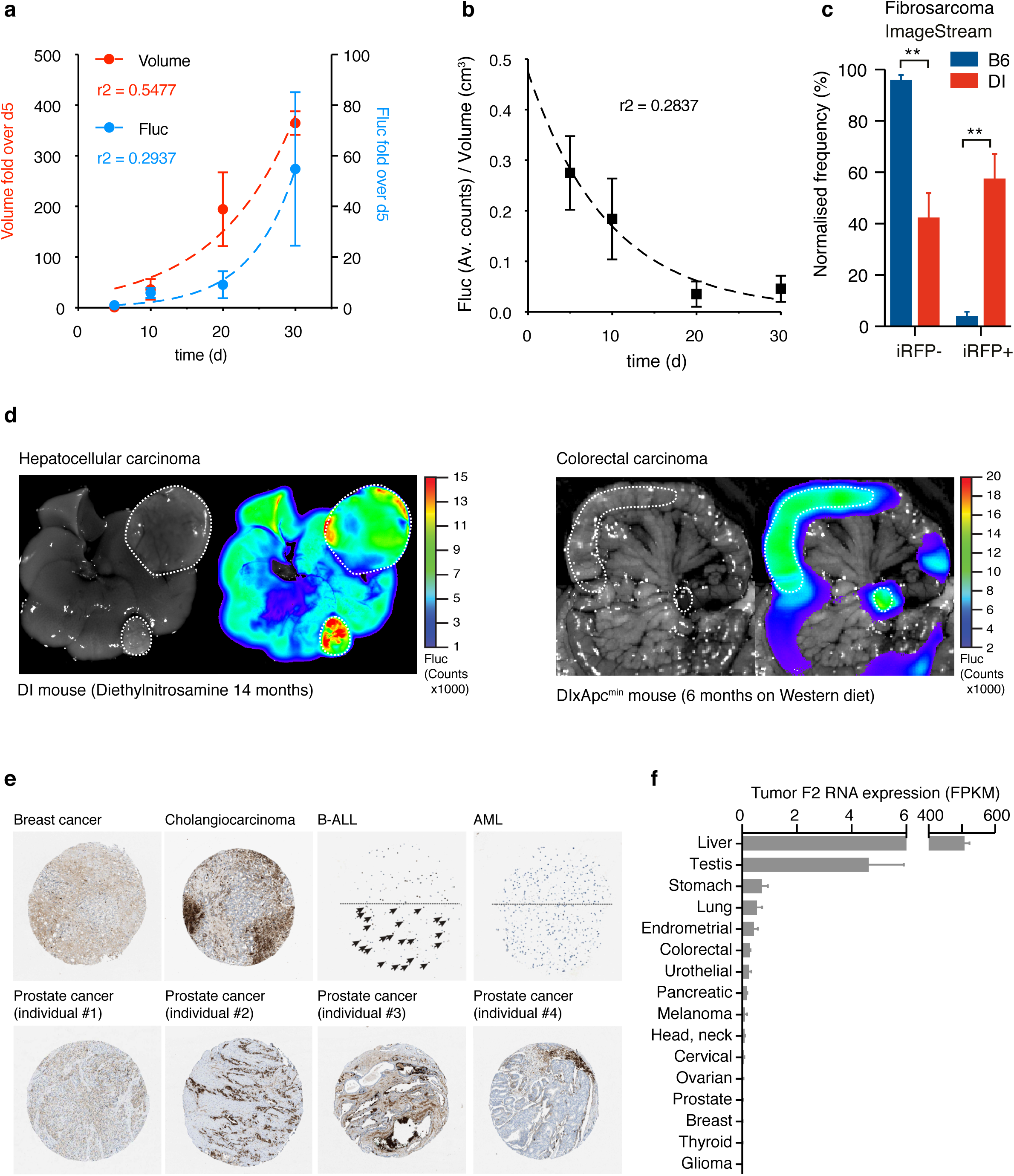
Tumor-derived prothrombin expression in mice and man. **a**, Increase of prothrombin expression (Fluc positivity) in D-Insight tumors transplanted in B6 animals. Non-linear regression (Exponential growth equation) indicated by dashed line. **b**, As rapidly growing tumors typically display areas of central necrosis, the relative Fluc signal normalized to tumor volume drops with time. Non-linear regressions (One phase decay) indicated by dashed line. **c**, ImageStream based confirmation of iRFP positivity of viable primary tumor cells obtained from D-Insight reporter positive fibrosarcomas compared to B6 control fibrosarcomas. **d**, Prothrombin positivity in murine hepatocellular and colorectal carcinoma (D-Insight animals treated with diethylnitrosamine for 14 months, and D-Insight animals on a Apc^min^ background after 6 months on Western diet, respectively). **e**, Prothrombin immunohistochemistry and F2 RNA seq data of human tumors, displaying various extent of positivity within and across tumor entities and individuals (data obtained from protein atlas and the National Cancer Institute’s Genomic Data Commons Data Portal (https://portal.gdc.cancer.gov)).

**Extended Data Figure 5.**
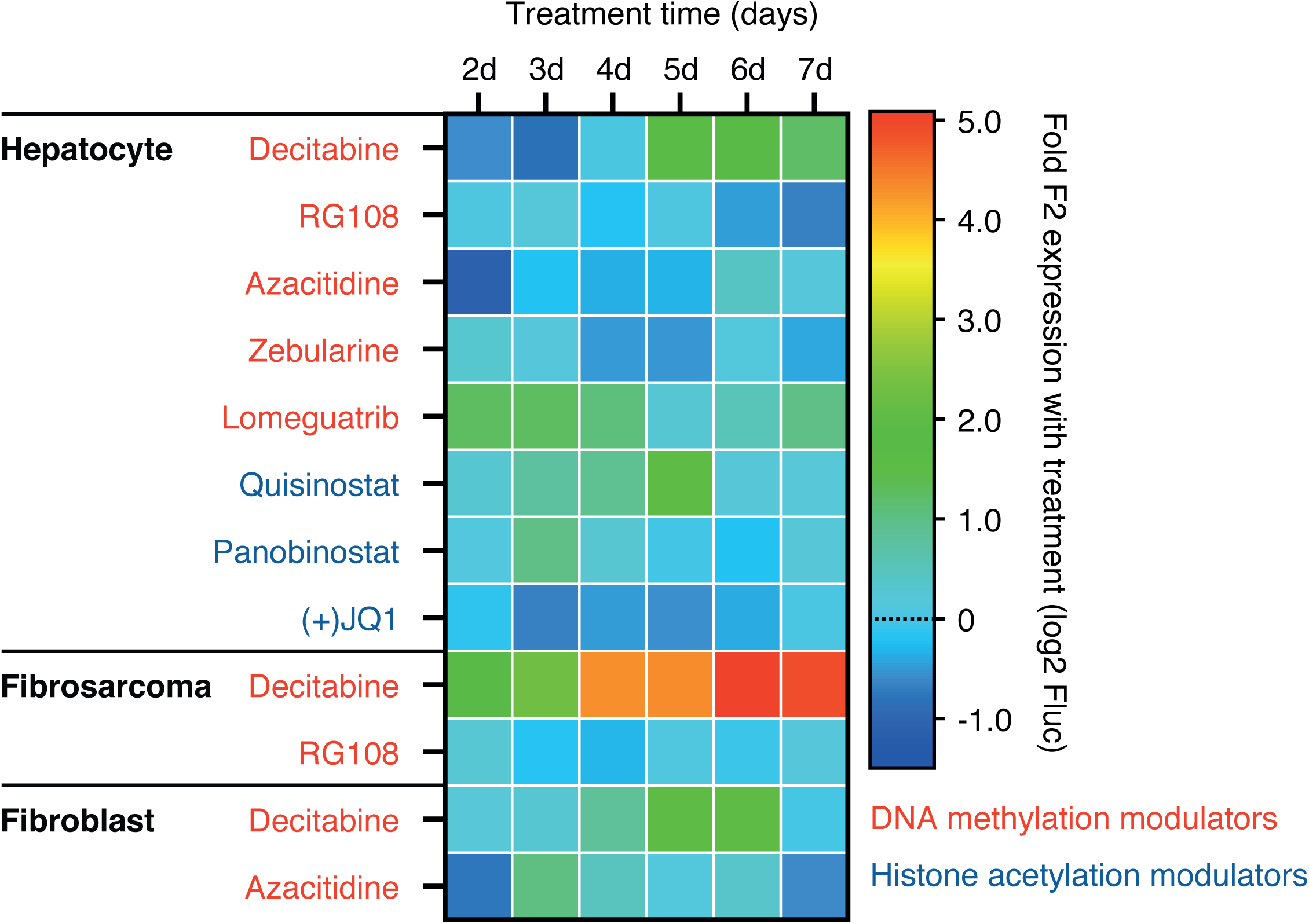
Tissue specific targeting of prothrombin gene expression. Small scale proof-of-concept study to illustrate that D-Insight derived primary cell cultures (hepatocytes, fibrosarcomas and fibroblasts) can be used to define tissue-specific therapeutic vulnerabilities.

**Supplementary Figure S1.**
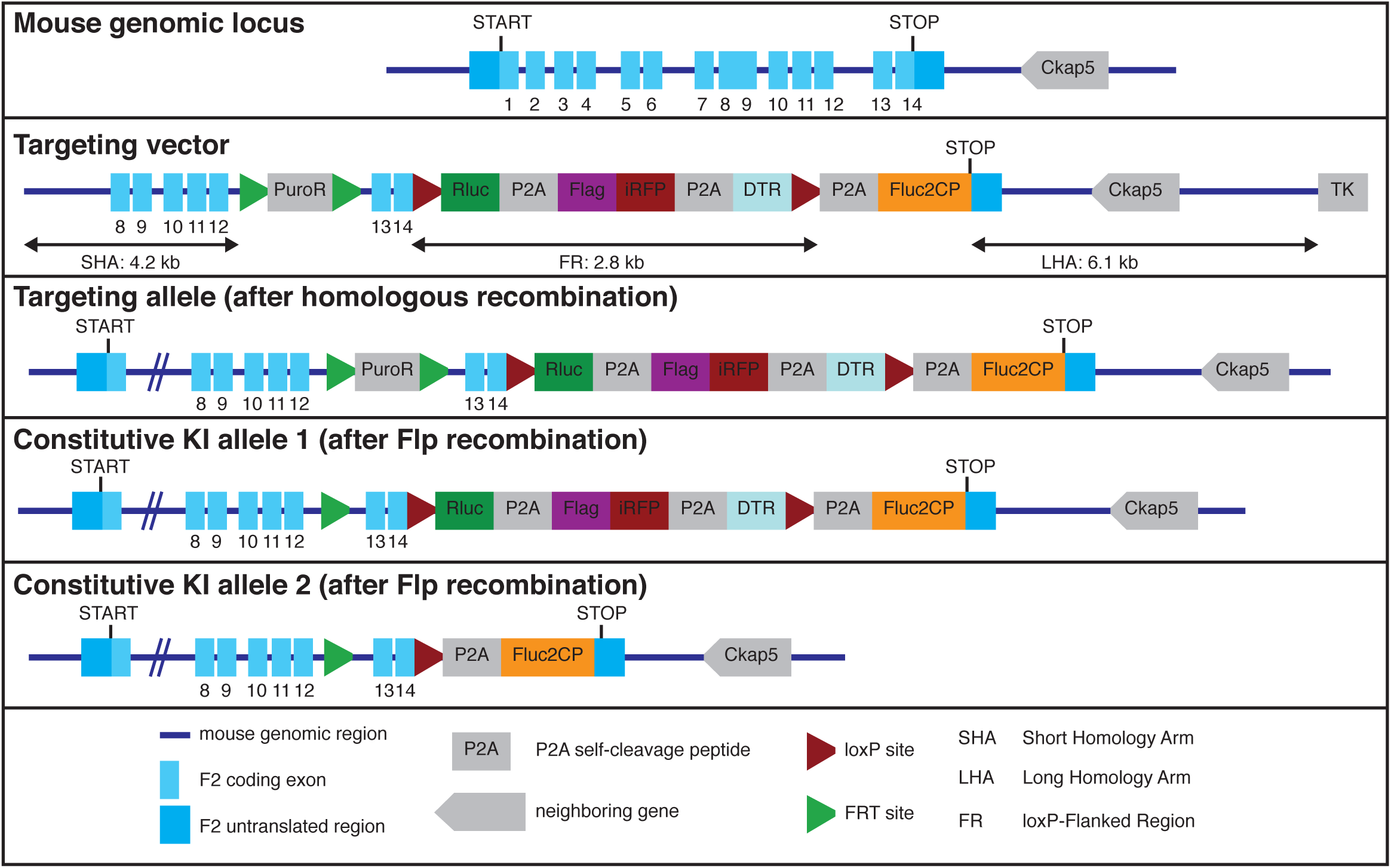

## References

1 Kim, M. S. et al. A draft map of the human proteome. Nature 509, 575–581, doi:10.1038/nature13302 (2014).

2 Uhlén, M. et al. Proteomics. Tissue-based map of the human proteome. Science 347, 1260419, doi:10.1126/science.1260419 (2015).

3 Rothman, J. E. & Orci, L. Molecular dissection of the secretory pathway. Nature 355, 409–415, doi:10.1038/355409a0 (1992).

4 Kelly, R. B. Pathways of protein secretion in eukaryotes. Science 230, 25–32, doi:10.1126/science.2994224 (1985).

5 Alberts, B. et al. Molecular Biology of the Cell. Garland Science, 2015.

6 Doroudgar, S. & Glembotski, C. C. The cardiokine story unfolds: ischemic stress-induced protein secretion in the heart. Trends Mol Med 17, 207–214, doi:10.1016/j.molmed.2010.12.003 (2011).

7 Egerstedt, A. et al. Profiling of the plasma proteome across different stages of human heart failure. Nat Commun 10, 5830, doi:10.1038/s41467-019-13306-y (2019).

8 Park-Windhol, C. & D’Amore, P. A. Disorders of Vascular Permeability. Annu Rev Pathol 11, 251–281, doi:10.1146/annurev-pathol-012615-044506 (2016).

9 Kuznetsov, G. & Nigam, S. K. Folding of secretory and membrane proteins. N Engl J Med 339, 1688–1695, doi:10.1056/nejm199812033392307 (1998).

10 Wang, M. & Kaufman, R. J. Protein misfolding in the endoplasmic reticulum as a conduit to human disease. Nature 529, 326–335, doi:10.1038/nature17041 (2016).

11 De Matteis, M. A. & Luini, A. Mendelian disorders of membrane trafficking. N Engl J Med 365, 927–938, doi:10.1056/NEJMra0910494 (2011).

12 Willis, M. S. & Patterson, C. Proteotoxicity and cardiac dysfunction--Alzheimer’s disease of the heart? N Engl J Med 368, 455–464, doi:10.1056/NEJMra1106180 (2013).

13 Carrell, R. W. & Lomas, D. A. Alpha1-antitrypsin deficiency--a model for conformational diseases. N Engl J Med 346, 45–53, doi:10.1056/NEJMra010772 (2002).

14 Tripodi, A. & Mannucci, P. M. The coagulopathy of chronic liver disease. N Engl J Med 365, 147–156, doi:10.1056/NEJMra1011170 (2011).

15 Schulze, R. J., Schott, M. B., Casey, C. A., Tuma, P. L. & McNiven, M. A. The cell biology of the hepatocyte: A membrane trafficking machine. J Cell Biol 218, 2096–2112, doi:10.1083/jcb.201903090 (2019).

16 Bernal, W. & Wendon, J. Acute liver failure. N Engl J Med 369, 2525–2534, doi:10.1056/NEJMra1208937 (2013).

17 Lozano, R. et al. Global and regional mortality from 235 causes of death for 20 age groups in 1990 and 2010: a systematic analysis for the Global Burden of Disease Study 2010. Lancet 380, 2095–2128, doi:10.1016/s0140-6736(12)61728-0 (2012).

18 Karamyshev, A. L., Tikhonova, E. B. & Karamysheva, Z. N. Translational Control of Secretory Proteins in Health and Disease. Int J Mol Sci 21, doi:10.3390/ijms21072538 (2020).

19 Danckwardt, S. et al. p38 MAPK controls prothrombin expression by regulated RNA 3’ end processing. Mol Cell 41, 298–310, doi:10.1016/j.molcel.2010.12.032 (2011).

20 Furukawa, T. et al. Angiogenic factor. Nature 356, 668, doi:10.1038/356668a0 (1992).

21 Radisky, D. C., Stallings-Mann, M., Hirai, Y. & Bissell, M. J. Single proteins might have dual but related functions in intracellular and extracellular microenvironments. Nat Rev Mol Cell Biol 10, 228–234, doi:10.1038/nrm2633 (2009).

22 Mani, M. et al. MoonProt: a database for proteins that are known to moonlight. Nucleic Acids Res 43, D277–282, doi:10.1093/nar/gku954 (2015).

23 Kim, J., Gee, H. Y. & Lee, M. G. Unconventional protein secretion - new insights into the pathogenesis and therapeutic targets of human diseases. J Cell Sci 131, doi:10.1242/jcs.213686 (2018).

24 Massoud, T. F. & Gambhir, S. S. Integrating noninvasive molecular imaging into molecular medicine: an evolving paradigm. Trends Mol Med 13, 183–191, doi:10.1016/j.molmed.2007.03.003 (2007).

25 Weissleder, R., Nahrendorf, M. & Pittet, M. J. Imaging macrophages with nanoparticles. Nat Mater 13, 125–138, doi:10.1038/nmat3780 (2014).

26 Farhadi, A., Sigmund, F., Westmeyer, G. G. & Shapiro, M. G. Genetically encodable materials for non-invasive biological imaging. Nat Mater 20, 585–592, doi:10.1038/s41563-020-00883-3 (2021).

27 van Dam, G. M. et al. Intraoperative tumor-specific fluorescence imaging in ovarian cancer by folate receptor-α targeting: first in-human results. Nat Med 17, 1315–1319, doi:10.1038/nm.2472 (2011).

28 Kvon, E. Z. Using transgenic reporter assays to functionally characterize enhancers in animals. Genomics 106, 185–192, doi:10.1016/j.ygeno.2015.06.007 (2015).

29 Close, D. M., Xu, T., Sayler, G. S. & Ripp, S. In vivo bioluminescent imaging (BLI): noninvasive visualization and interrogation of biological processes in living animals. Sensors (Basel) 11, 180–206, doi:10.3390/s110100180 (2011).

30 Reik, W. Stability and flexibility of epigenetic gene regulation in mammalian development. Nature 447, 425–432, doi:10.1038/nature05918 (2007).

31 Manning, K. S. & Cooper, T. A. The roles of RNA processing in translating genotype to phenotype. Nat Rev Mol Cell Biol 18, 102–114, doi:10.1038/nrm.2016.139 (2017).

32 Esteller, M. Non-coding RNAs in human disease. Nat Rev Genet 12, 861–874, doi:10.1038/nrg3074 (2011).

33 Di Cera, E. Thrombin. Mol Aspects Med 29, 203–254, doi:10.1016/j.mam.2008.01.001 (2008).

34 Di Cera, E. Thrombin interactions. Chest 124, 11s–17s, doi:10.1378/chest.124.3_suppl.11s (2003).

35 Borissoff, J. I., Spronk, H. M. & ten Cate, H. The hemostatic system as a modulator of atherosclerosis. N Engl J Med 364, 1746–1760, doi:10.1056/NEJMra1011670 (2011).

36 Danckwardt, S., Hentze, M. W. & Kulozik, A. E. Pathologies at the nexus of blood coagulation and inflammation: thrombin in hemostasis, cancer, and beyond. J Mol Med (Berl) 91, 1257–1271, doi:10.1007/s00109-013-1074-5 (2013).

37 Coughlin, S. R. Thrombin signalling and protease-activated receptors. Nature 407, 258–264, doi:10.1038/35025229 (2000).

38 Loening, A. M., Wu, A. M. & Gambhir, S. S. Red-shifted Renilla reniformis luciferase variants for imaging in living subjects. Nat Methods 4, 641–643, doi:10.1038/nmeth1070 (2007).

39 Filonov, G. S. et al. Bright and stable near-infrared fluorescent protein for in vivo imaging. Nat Biotechnol 29, 757–761, doi:10.1038/nbt.1918 (2011).

40 Kim, J. H. et al. High cleavage efficiency of a 2A peptide derived from porcine teschovirus-1 in human cell lines, zebrafish and mice. PLoS One 6, e18556, doi:10.1371/journal.pone.0018556 (2011).

41 Szymczak, A. L. et al. Correction of multi-gene deficiency in vivo using a single ‘self-cleaving’ 2A peptide-based retroviral vector. Nat Biotechnol 22, 589–594, doi:10.1038/nbt957 (2004).

42 Dragulescu-Andrasi, A., Chan, C. T., De, A., Massoud, T. F. & Gambhir, S. S. Bioluminescence resonance energy transfer (BRET) imaging of protein-protein interactions within deep tissues of living subjects. Proc Natl Acad Sci U S A 108, 12060–12065, doi:10.1073/pnas.1100923108 (2011).

43 Xiong, L., Shuhendler, A. J. & Rao, J. Self-luminescing BRET-FRET near-infrared dots for in vivo lymph-node mapping and tumour imaging. Nat Commun 3, 1193, doi:10.1038/ncomms2197 (2012).

44 Saito, M. et al. Diphtheria toxin receptor-mediated conditional and targeted cell ablation in transgenic mice. Nat Biotechnol 19, 746–750, doi:10.1038/90795 (2001).

45 Sauer, B. & Henderson, N. Site-specific DNA recombination in mammalian cells by the Cre recombinase of bacteriophage P1. Proc Natl Acad Sci U S A 85, 5166–5170, doi:10.1073/pnas.85.14.5166 (1988).

46 Soifer, S. J., Peters, K. G., O’Keefe, J. & Coughlin, S. R. Disparate temporal expression of the prothrombin and thrombin receptor genes during mouse development. Am J Pathol 144, 60–69 (1994).

47 Dihanich, M., Kaser, M., Reinhard, E., Cunningham, D. & Monard, D. Prothrombin mRNA is expressed by cells of the nervous system. Neuron 6, 575–581, doi:10.1016/0896-6273(91)90060-d (1991).

48 Munns, T. W., Johnston, M. F., Liszewski, M. K. & Olson, R. E. Vitamin K-dependent synthesis and modification of precursor prothrombin in cultured H-35 hepatoma cells. Proc Natl Acad Sci U S A 73, 2803–2807, doi:10.1073/pnas.73.8.2803 (1976).

49 Sun, W. Y. et al. Prothrombin deficiency results in embryonic and neonatal lethality in mice. Proc Natl Acad Sci U S A 95, 7597–7602, doi:10.1073/pnas.95.13.7597 (1998).

50 Xue, J. et al. Incomplete embryonic lethality and fatal neonatal hemorrhage caused by prothrombin deficiency in mice. Proc Natl Acad Sci U S A 95, 7603–7607, doi:10.1073/pnas.95.13.7603 (1998).

51 Welch, E. M. et al. PTC124 targets genetic disorders caused by nonsense mutations. Nature 447, 87–91, doi:10.1038/nature05756 (2007).

52 Hengstler, J. G. et al. Cultures with cryopreserved hepatocytes: applicability for studies of enzyme induction. Chem Biol Interact 125, 51–73, doi:10.1016/s0009-2797(99)00141-6 (2000).

53 Khorana, A. A., Francis, C. W., Culakova, E., Kuderer, N. M. & Lyman, G. H. Thromboembolism is a leading cause of death in cancer patients receiving outpatient chemotherapy. J Thromb Haemost 5, 632–634, doi:10.1111/j.1538-7836.2007.02374.x (2007).

54 Timp, J. F., Braekkan, S. K., Versteeg, H. H. & Cannegieter, S. C. Epidemiology of cancer-associated venous thrombosis. Blood 122, 1712–1723, doi:10.1182/blood-2013-04-460121 (2013).

55 Noy, R. & Pollard, J. W. Tumor-associated macrophages: from mechanisms to therapy. Immunity 41, 49–61, doi:10.1016/j.immuni.2014.06.010 (2014).

56 Xue, Y. H. et al. Thrombin is a therapeutic target for metastatic osteopontin-positive hepatocellular carcinoma. Hepatology 52, 2012–2022, doi:10.1002/hep.23942 (2010).

57 Karpatkin, S., Finlay, T. H., Ballesteros, A. L. & Karpatkin, M. Effect of warfarin on prothrombin synthesis and secretion in human Hep G2 cells. Blood 70, 773–778 (1987).

58 Hisada, Y. & Mackman, N. Cancer-associated pathways and biomarkers of venous thrombosis. Blood 130, 1499–1506, doi:10.1182/blood-2017-03-743211 (2017).

59 Rickles, F. R. & Falanga, A. Molecular basis for the relationship between thrombosis and cancer. Thromb Res 102, V215–224, doi:10.1016/s0049-3848(01)00285-7 (2001).

60 Sonawane, A. R. et al. Understanding Tissue-Specific Gene Regulation. Cell Rep 21, 1077–1088, doi:10.1016/j.celrep.2017.10.001 (2017).

61 Oh, P. et al. Subtractive proteomic mapping of the endothelial surface in lung and solid tumours for tissue-specific therapy. Nature 429, 629–635, doi:10.1038/nature02580 (2004).

62 Rabouille, C. Pathways of Unconventional Protein Secretion. Trends Cell Biol 27, 230–240, doi:10.1016/j.tcb.2016.11.007 (2017).

63 Zhang, M. et al. A Translocation Pathway for Vesicle-Mediated Unconventional Protein Secretion. Cell 181, 637-652.e615, doi:10.1016/j.cell.2020.03.031 (2020).

64 Krenzlin, H., Lorenz, V., Danckwardt, S., Kempski, O. & Alessandri, B. The Importance of Thrombin in Cerebral Injury and Disease. Int J Mol Sci 17, doi:10.3390/ijms17010084 (2016).

65 Poort, S. R., Rosendaal, F. R., Reitsma, P. H. & Bertina, R. M. A common genetic variation in the 3’-untranslated region of the prothrombin gene is associated with elevated plasma prothrombin levels and an increase in venous thrombosis. Blood 88, 3698–3703 (1996).

66 Danckwardt, S. et al. The prothrombin 3’end formation signal reveals a unique architecture that is sensitive to thrombophilic gain-of-function mutations. Blood 104, 428–435, doi:10.1182/blood-2003-08-2894 (2004).

67 Kopec, A. K. et al. Thrombin promotes diet-induced obesity through fibrin-driven inflammation. J Clin Invest 127, 3152–3166, doi:10.1172/jci92744 (2017).

68 Postic, C. et al. Dual roles for glucokinase in glucose homeostasis as determined by liver and pancreatic beta cell-specific gene knock-outs using Cre recombinase. J Biol Chem 274, 305–315, doi:10.1074/jbc.274.1.305 (1999).

69 Ogorodnikov, A. et al. Transcriptome 3’end organization by PCF11 links alternative polyadenylation to formation and neuronal differentiation of neuroblastoma. Nat Commun 9, 5331, doi: 10.1038/s41467-018-07580-5 (2018).

